# Structural insight into molecular inhibitory mechanism of InsP_6_ on African Swine Fever Virus mRNA-decapping enzyme g5Rp

**DOI:** 10.1101/2021.06.23.449648

**Authors:** Yan Yang, Changhui Zhang, Li Li, Xuehui Li, Xin Yang, Yao Zhao, Cheng Chen, Wei Wang, Zhihui Zhong, Cheng Yang, Zhen Huang, Dan Su

**Affiliations:** State Key Laboratory of Biotherapy and Cancer Center, West China Hospital, Sichuan University and Collaborative Innovation Center of Biotherapy, Chengdu, Sichuan 610041, P.R. China; Institute of Life Sciences, Chongqing Medical University, Chongqing 400016, P.R. China; Shanghai Institute for Advanced Immunochemical Studies and School of Life Science and Technology, ShanghaiTech University, Shanghai, China; School of Life Sciences, Tianjin University, Tianjin 300072, P.R. China; College of Life Sciences, Sichuan University, Chengdu, 610041, China; Tianjin International Joint Academy of Biotechnology and Medicine, Tianjin 300457, P.R. China

**Author notes:** Address correspondence to Dan Su, **Email**.

**Keywords:** g5Rp, ASFV, mRNA decapping enzyme, Nudix hydrolases, InsP_6_

## Abstract

Removal of 5′ cap on cellular mRNAs by the African Swine Fever Virus (ASFV) decapping enzyme g5R protein (g5Rp) is beneficial to viral gene expression during the early stages of infection. As the only nucleoside diphosphate linked moiety X (Nudix) decapping enzyme encoded in the ASFV genome, g5Rp works in both the degradation of cellular mRNA and hydrolyzation of the diphosphoinositol polyphosphates. Here, we report the structures of dimeric g5Rp and its complex with inositol hexakisphosphate (InsP_6_). The two g5Rp protomers interact head-to-head to form a dimer, and the dimeric interface is formed by extensive polar and nonpolar interactions. Each protomer composed a unique N-terminal helical domain and C-terminal classic Nudix domain. As a mRNA decapping enzyme, we identified key residues, including K^8^, K^94^, K^95^, K^98^, K^175^, R^221^, and K^243^ located on the substrate RNA binding interfaces of g5Rp, are important to RNA binding and decapping enzyme activity. Furthermore, we identified that the g5Rp-mediated mRNA decapping was inhibited by the InsP_6_. The g5Rp–InsP_6_ complex structure showed that the InsP_6_ molecules occupy the same regions that primarily mediate g5Rp-RNA interaction, elucidating the roles of InsP_6_ in the regulation of the viral decapping activity of g5Rp in mRNA degradation. Collectively, these results provide the structural basis of interaction between RNA and g5Rp and highlight the inhibitory mechanism of InsP_6_ on mRNA decapping by g5Rp.

## Introduction

African swine fever virus (ASFV), which is an enveloped double-stranded DNA virus, has a genome that varies between 170 and 193 kilobase pairs with 151–167 open reading frames depending on the virus strain ^1,2^. As the only known DNA arbovirus, ASFV is the sole member of the *Asfarviridae*, a family of African swine fever-like viruses that are relatively independent of the host cell transcriptional machinery for viral replication ^3,4^. The ASFV infection of domestic swine can result in various disease forms, ranging from highly lethal to subclinical depending on the contributing viral and host factors ^5^. Since 2018, ASFV has spread into China and led to a high mortality rate in domestic pigs ^6,7^. Currently, there are still no effective vaccines or specific drugs available against this particular virus ^8,9^.

During an ASFV infection, protein synthesis in the host cell is inhibited as a result of a massive degradation of host cellular mRNAs in the cytoplasm of infected cells ^10,11^. As part of its strategy to inhibit host cellular translation and promote viral protein synthesis instead, the virus targets the mRNAs of the host cell using specific enzymes^12^. Hydrolysis of the 5′ cap structure (m^7^GpppN) on eukaryotic mRNAs, a process known as decapping, is considered to be a crucial and highly regulated step in the degradation of mRNA^13^. Some viruses including ASFV and vaccinia virus (VACV) can harbor decapping enzymes for control of viral and cellular gene expression^14^. Two poxvirus Nudix hydrolases D9, D10 have been confirmed with intrinsic mRNA-decapping activity, although the two decapping enzymes appear to have some differences in substrate recognition ^15,16^.

Nudix hydrolases (nucleoside diphosphate-linked moiety X) are widely present in bacteria, archaea, and eukarya, where they belong to a superfamily of hydrolytic enzymes that catalyze the cleavage of nucleoside diphosphates and the decapping of the 5′ cap of mRNAs, the latter of which plays a pivotal role in mRNA metabolism ^17,18^. Mammalian cells have about 30 different genes with Nudix motifs, including Dcp2, Nudt16, and NUDT3/DIPP1, which cleaves mRNA caps in mRNA degradation by the 5′-3′ decay pathway *in vivo*^19–21^. The mRNA-decapping enzyme g5Rp, which is the only Nudix hydrolase in ASFV, shares sequence similarity to the mRNA-decapping enzymes Dcp2 in *Schizosaccharomyces pombe* and D9 or D10 in the VACV ^22–24^. However, g5Rp and its Nudix homologs D9 and D10 exhibit higher hydrolytic activity toward diphosphoinositol polyphosphates and dinucleotide polyphosphates than toward cap analogs ^25,26^. Similar to Dcp2, these Nudix hydrolases cleave the mRNA cap attached to an RNA moiety predicating that RNA binding is crucial for performing its mRNA decapping activity^16^. Recently, structural study has confirmed that the Nudix protein CFI_m_25 has a sequence-specific RNA-binding capability ^27^. The requirement of RNA binding for majority of the Nudix decapping enzymes suggest that the members of Nudix family also belong to RNA-binding proteins.

The viral mRNA-decapping enzyme g5Rp is expressed in the endoplasmic reticulum from the early stage of ASFV infection and accumulates throughout the infection process, playing an essential role in the machinery assembly of mRNA regulation and translation initiation ^23^. Likes other members of Nudix family, g5Rp has a broader range of nucleotide substrate specificity, including that for a variety of guanine and adenine nucleotides and dinucleotide polyphosphates^25^. Generally, g5Rp has two distinct enzymatic activities *in vitro* (viz., diadenosine hexaphosphate hydrolase activity and mRNA-decapping activity), implying that it plays roles in viral membrane morphogenesis and mRNA regulation during viral infections ^28^. In light of these biochemical observations, the elucidation of the structure of g5Rp is of fundamental importance toward our understanding of the molecular mechanisms through which it degrades cellular RNAs and regulates viral gene expression.

Here, we report the crystal structure of g5Rp and its complex structure with InsP_6_. Combing with biochemical experiments, the dimeric form of g5Rp and three RNA binding surfaces on each protomer are critical to substrate RNA binding of g5Rp. The g5Rp-InsP_6_ complex structure shows that two of RNA binding surfaces are occupied by InsP_6_ indicating that InsP_6_ may play a role in its ability to inhibit g5Rp-RNA binding activity. Meanwhile, we evaluate the inhibitor effect of InsP_6_ on the mRNA decapping enzyme activity of g5Rp. Therefore, we proposed that such inhibition could be caused by the competition of InsP_6_ with substrate mRNA for binding to the g5Rp. Furthermore, we show detail how InsP_6_ inhibits g5Rp activity by occupied the RNA-binding interfaces on g5Rp, thereby competitively blocking the binding of substrate mRNA to the enzyme. These results suggest InsP_6_ or its structural analogs may be involved in the manipulation of the mRNA decapping process during viral infections and provide an essential structural basis for the development of ASFV chemotherapies in the future.

## Results

### Characterization of recombinant ASFV g5Rp protein

Recombinant wild type ASFV g5Rp (residues 1–250) was expressed in *Escherichia coli* (*E. coli*) with a N-terminal His_6_-tag. The purified g5Rp protein was eluted from a Superdex 200 column (GE Healthcare) with a major elution volume of 15.6 mL, indicating an approximate molecular weight of 32.1 kDa (Fig. S1*A*). The fractions were further analyzed by sodium dodecyl sulfate polyacrylamide gel electrophoresis (SDS/PAGE), showing a g5Rp band of 29.9 kDa (Fig. S1*B*). Crosslinking assay confirmed that the g5Rp protein exits as a stable homodimer in solution (Fig. S1*C*).

We first characterized the nucleic acid binding ability of g5Rp with different length of single-stranded RNA (12-mer, 26-mer ssRNA). Electrophoretic Mobility Shift Assays (EMSAs) results demonstrated that g5Rp binds ssRNA (0.25 μM) at the lowest concentration of 0.5 μM (Fig. 1*A* and *B*). Furthermore, we measured the binding affinity of wild type (WT) g5Rp for ssRNA by using surface plasmon resonance (SPR). (Fig. 1*D* and *E*). The enzyme exhibited a stronger binding affinity to ssRNAs with equilibrium dissociation constant 12-mer *K_D_*= 164.0 nM, 26-mer *K_D_* = 44.8 nM. The kinetic analysis of the binding experiments is shown in Supplementary Table 1. These results indicate that g5Rp possesses a higher affinity with long ssRNA. Next, we re-evaluated the decapping activity of recombinant g5Rp by incubating the protein with a ^32^P-cap-labeled RNA substrate in a reaction. The products of the reaction were resolved by polyethyleneimine (PEI)-cellulose thin layer chromatography (TLC) and detected by autoradiography^23^. As shown in Fig 1*C*, the recombinant g5Rp in the decapping reaction released 7-methylguanosine cap (m^7^GDP) product efficiently. On the contrary, the ^32^P-cap-labeled RNA substrate as control remained at the origin of the plate. These results suggest that the recombinant g5Rp possesses efficient mRNA decapping enzyme activity.

**FIG 1.**
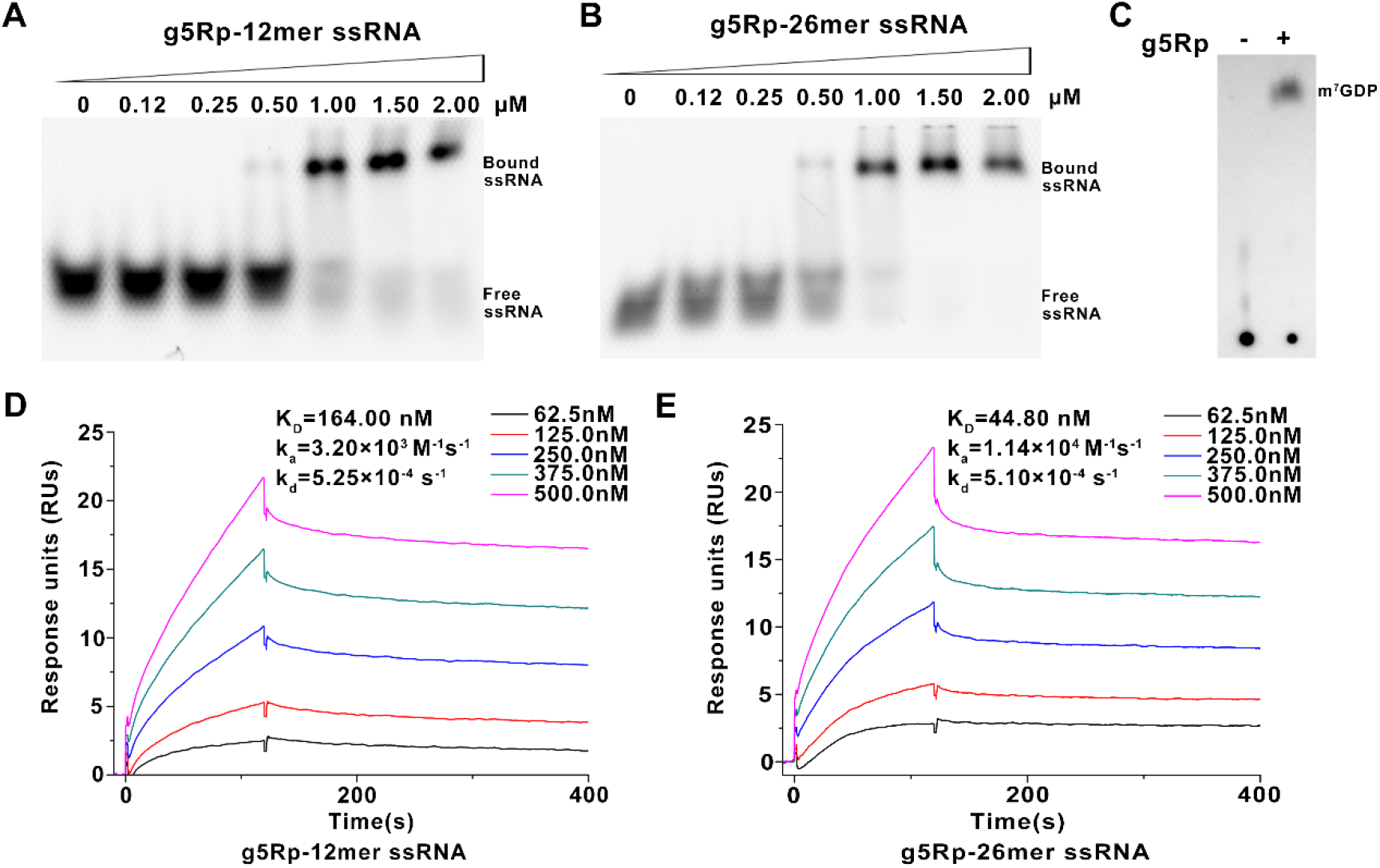
Characterization of the African swine fever virus decapping enzyme g5Rp. **(A-B**) The binding abilities of g5Rp to 12-mer and 26-mer ssRNA were determined by EMSA, in the reaction, the protein concentration is marked on the figure, and the nucleic acid concentration is 0.25uM. (**C**) The decapping activity of g5Rp. (**D-E**) The binding abilities of g5Rp to 12-mer and 26-mer ssRNA were determined by surface plasmon resonance.

### Overview of the ASFV g5Rp structure

To investigate structural insights into the catalytic mechanism of g5Rp, we determined its dimeric structure by single-wavelength anomalous diffraction (SAD) phases using selenomethionine-labeled protein (supplementary result). As shown in Fig. 2*A*, the g5Rp dimer composed two protomers that each adopts a baseball glove shape with distinct N-terminal helical domain and C-terminal Nudix domain (Fig. 2*B*). The helical domain (residues 36–124) forms a globin-fold-like feature composed of six α-helices (α1 –α6) that connect to the Nudix domain by two hinge linkers (linker I: residues 32–35; linker II: residues 119–139). The Nudix domain (residues 1–35 and 125–250) consists of a central curved β-sheet (β1, β2, β3, β4) surrounded by five α-helices (α7–α11) and several loops, thereby forming a classic α–β–α sandwich structure. Linker II splits the top of the β-sheet to connect α6 and α7 together (Fig. 2*C*). The Nudix motif located in center of the Nudix domain is a highly conserved comprising the loop-helix-loop architecture that contains the Nudix signature sequence extending from residues ^132^GKPKEDESDLTCAIREFEEETGI^154^ in g5Rp (Fig. 2*D*). The sequence of the g5Rp Nudix motif matches the classic pattern of the Nudix motif in the Nudix hydrolase superfamily; that is, GX5EX7REUXEEXGU, where X is any residue and U is Ile, Leu, or Val ^29,30^. Using the Dali server^31^, we compared the structure of the g5Rp with that of other proteins in the Protein Data Bank (PDB), whereupon 46 structures were found to be likely homologous to the enzyme, with Z-scores in the range of 8–20 (Supplementary, Table 2). However, all the listed protein structures only shared high architectural similarity with the Nudix domain of g5Rp. Therefore, a search on the Dali server was carried out for the N-terminal domain alone, whereupon no homologous structure with a Z-score above 4 was found, suggesting that the N-terminus of g5Rp adopts a novel fold. Comparing with the structures of Dcp2 in a number of different conformations, the g5Rp shows a unique N-terminal globin-fold-like domain (Fig. S1*D* and *E*).

**FIG 2.**
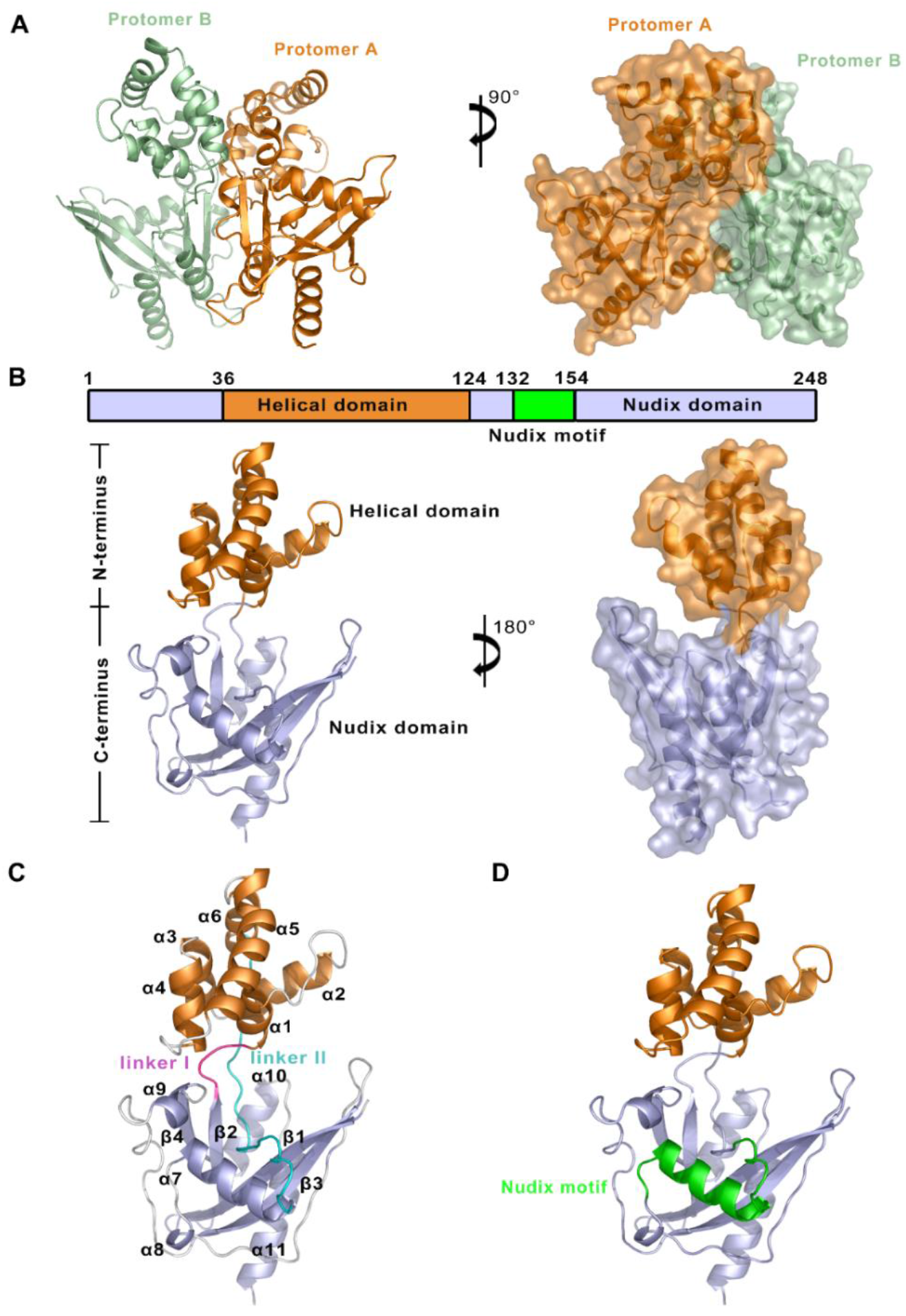
Structure of the African swine fever virus decapping enzyme g5Rp. (**A**) The dimeric structure of the African swine fever virus decapping enzyme g5Rp. The dimer consists of protomers A and B in a back-to-back orientation. Protomer A is colored orange and protomer B is colored pale green. (**B**) “Boxing glove” arrangement of the overall structure of g5Rp. The division of the domain is based on the g5Rp structure, where each domain is color coded, with the helical domain in orange and the Nudix domain in light blue (containing Nudix motif, in green). The N-terminus and C-terminus domains are marked by black box. (**C**) The details structure information of g5Rp. The 11 α-helices, 4 β-sheets, and loops in the structure. The loops are colored in white, except for linker I (in magenta) and linker II (in cyan). (**D**) The details of Nudix motif in g5Rp. The Nudix motif is colored by green.

A previously study showed that the N-terminus of g5Rp is the major mediator of RNA interaction ^28^. However, the positively charged surface of g5Rp structure overlaps both the N- and C-terminal region of protein that may exhibit RNA-binding activity (Fig. S2*A*). We proposed that both positively charged regions could contribute to g5Rp-RNA interaction. To test the hypothesis, we measured the binding of the truncation variants g5RpΔC (helical domain, residues 36-124) and g5RpΔN (Nudix domain, connecting residues 1-35 and 125-250 directly) to single-stranded ssRNA (12-mer, 26-mer), respectively. Our EMSA results showed that both helical domain (g5RpΔC) and Nudix domain (g5RpΔN) of g5Rp are involved in ssRNA interaction (Fig. S2*B*). The helical domain exhibited *K_D_* values of 39.0 and 50.7 nM for the surface-immobilized 12- and 26- mer ssRNAs measured by SPR, respectively (Fig. S2*C* and *D*). By contrast, the *K_D_* values of g5Rp wild-type for the ssRNAs (12-mer *K_D_* = 164.0nM, 26-mer *K_D_* = 44.8nM) are slightly lower than that of the helical domain with ssRNAs, which indicating that both full-length and truncated g5Rp associated with RNA with high affinity.

### The dimeric structure of g5Rp

When recombinant g5Rp was subject to gel filtration chromatogram to estimate molecular weight, it migrated as a single population of molecules at a molecular mass consistent with a monomer. However, g5Rp dimerization was consistent with cross-linking experiments (Fig. S1*A* and *C*). To obtain more information about the interfaces and likely biological assemblies of g5Rp, we analyzed its structure using the PDB-related interactive tool Proteins, Interfaces, Structures and Assemblies (PDBePISA)^32^. The results suggested that the g5Rp forms a stable symmetric dimer in crystal packing. The stable dimer was composed of two protomers (A and B) positioned in an orientation similar to two baseball gloves stuck together back-to-back (Fig. 2*A*). The dimer interfaces were stabilized mainly by hydrophobic interactions. Furthermore, a network of hydrogen bonds conferred additional stability to the interface. One interface was composed of four α-helices (α3 and α4 from each A and B protomer) from the N-terminus of each protomer. Residues Ile^64^, Asn^65^, Arg^67^, Leu^68^, Leu^69^, Lys^71^, Thr^72^, Arg^77^, Tyr^80^, His^81^ and Ile^84^ located in helices α3 and α4 played pivotal roles in stabilizing the dimeric form of the protein. The other dimer interface was located at the linker II portion of the C-terminus, and one solvent contact surface of the C-terminal domain. Residues Ile^116^, Asn^117^, Ala^119^, Lys^120^, Gly^121^, Ser^122^, Gly^123^, and Thr^124^ located on linker II, and residues Asn^195^, Met^196^, Leu^198^, Ser^199^, Leu^200^, Gln^201^, Ile^206^, Ser^210^, Lys^211^, Gln^215^, Glu^218^ and Ala^219^ at the C-terminus, forming hydrogen bonds in the dimer interface, with further contributions from a hydrophobic patch composed of Ile^206^, Ile^209^, Phe^222^, and Ile^223^ (Fig. S3*A*). To determine the multimeric state of g5Rp in solution and to examine which of its termini is critical for its dimerization, we measured the multimerization of two g5Rp truncation variants (g5RpΔN and g5RpΔC) using cross-linking experiments. The results showed that the wild type, N-terminus, and C-terminus of g5Rp all formed a dimeric conformation in solution (Fig. S3*B* and *C*). The g5Rp mutant I84A/I116A/L200A/I206A/F222A that prepared to dissociate the dimeric form of g5Rp was successful to alter a monomeric state, even the dimeric total buried area of 3050 Å^2^. Wild-type g5Rp and mutants were subjected to gel-filtration chromatography resulting that the mutant I84A/I116A/L200A/I206A/F222A has a larger retention volume, corresponding to a smaller molecular weight (Fig. S4*A)*. The protein cross-linking experiment showed that the dimeric conformation was significantly reduced in solution of mutant (Fig. S4*B*). The ssRNA binding ability of monomeric mutant has been measured by SPR and EMSA. Various concentration of monomeric mutant were passed over immobilized ssRNA. The resultant sensorgarms are shown in (Fig. S4*C* and *D*), and kinetic analysis is shown in Supplementary Table 1. EMSA data is shown in Fig. S4*E*. Both measurements produced consistent results indicating that the g5Rp mutant I84A/I116A/L200A/I206A/F222A partially impaired the RNA binding ability. Therefore, we proposed that the dimeric g5Rp is preferred to efficient RNA-binding.

### Structure of the g5Rp-InsP_6_ complex

The g5Rp was originally characterized through its ability to dephosphorylate 5-PP-InsP5 (InsP_7_) to produce InsP_6_ ^25^. We were surprised to find a tight interaction between InsP_6_ and g5Rp by microscale thermophoresis (MST) (Fig. 3*A*). To gain insight into the molecular basis of the interaction, we determined the crystal structure of the g5Rp–InsP_6_ complex and found that each asymmetric unit contained one g5Rp-InsP_6_ complex in space group *P*4_1_22. PDBePISA analysis revealed that an identical dimeric conformation exists in g5Rp–InsP_6_ complex structure (Fig. 3*B*). Two InsP_6_ molecules were situated on the edge of the β1 strand of each g5Rp protomer through interactions with residues Gln^6^, Lys^8^, and Lys^133^ (Fig. 3*C*). Due to the two-fold symmetry in the crystal, each of the g5Rp protomers shared two InsP_6_ molecules (InsP_6_ and InsP_6_ ^asym^) with its neighboring g5Rp protomer in the crystal lattice. Besides the InsP_6_ binding on the β1 strand located on the edge of Nudix domain, an extra InsP_6_ molecule from the neighboring molecule also interacted with g5Rp through residues Lys^94^ and Lys^98^ on α5 helix in helical domain (Fig. 3*C*). In this way, each InsP_6_ molecule is surrounded by four Lys residues in complex structure. The solvent-accessible surface of the InsP_6_-binding region of g5Rp was calculated according to the electrostatic potential. It was apparent that both InsP_6_ molecules were situated on the highly positively charged area located in the protein cleft between helical domain and Nudix domain of g5Rp (Fig. S5*A*). The local conformational changes of g5Rp in the complex structure induced by its interaction with InsP_6_ are illustrated in Fig. S5*B*. In the complex structure, the β1, β3, and β5 strands located in the C-terminal region had moved closer to the helical domain, and α2 was pushed away from the InsP_6_-binding sites. These changes rendered the g5Rp conformation more stable in the complex.

**FIG 3.**
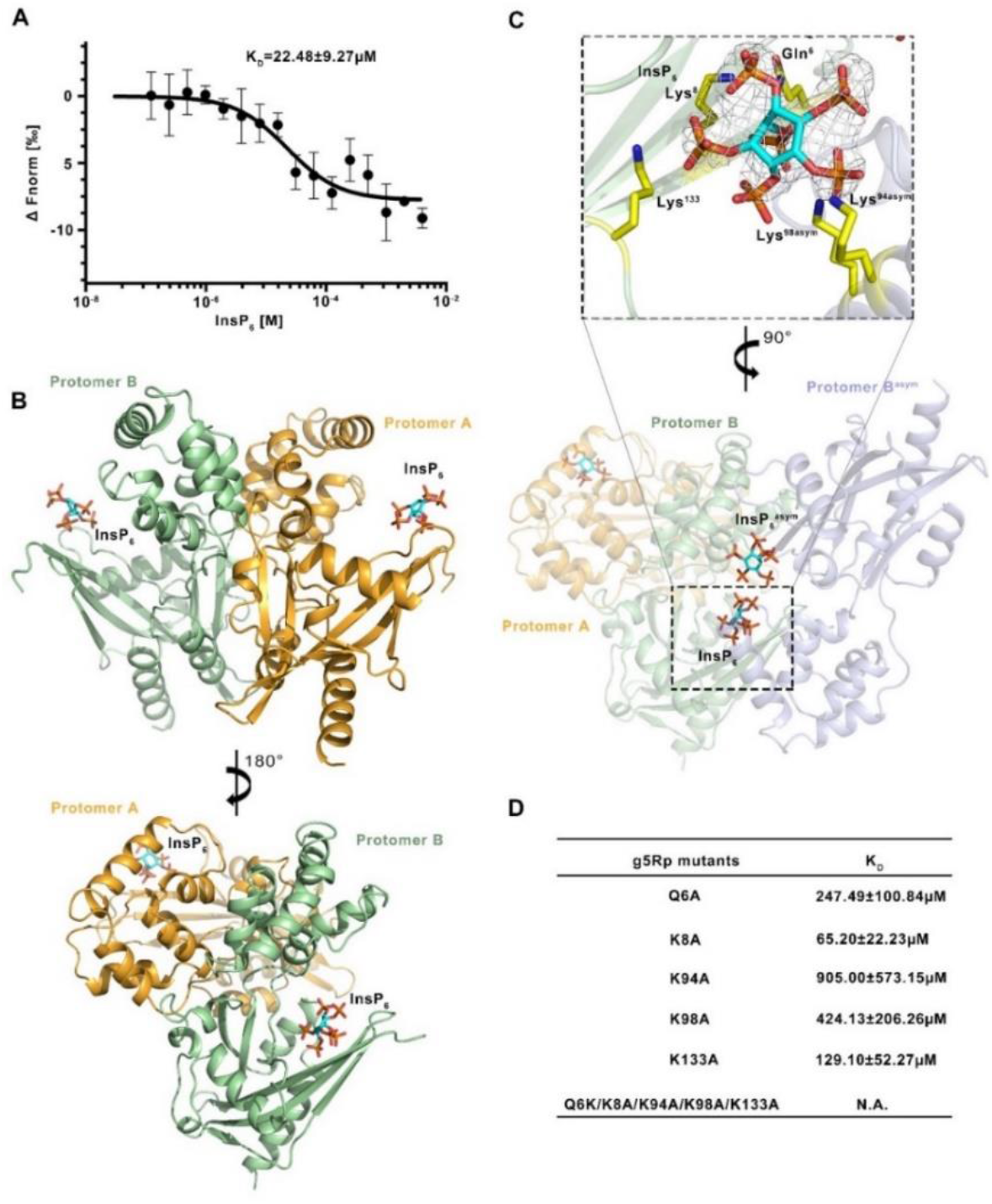
Structure of the g5Rp–InsP_6_ complex. (**A**) The binding ability between g5Rp with InsP_6_ is measured by MST. The dissociation constant was calculated from three independent replicates (shown as mean±s.d.) (**B**) The dimer of g5Rp in complex with InsP_6_. Details of g5Rp in complex with InsP_6_. The electron density of InsP_6_ is shown as the 2Fo–Fc map contoured at 0.8 σ and generated with InsP_6_ omitted. (**D**) The binding ability between various g5Rp mutants and InsP_6_ is measured by MST. The dissociation constant between g5Rp mutants with InsP_6_ were calculated from three independent replicates (shown as mean±s.d.). InsP_6_ is shown as a stick model.

To assess their relative importance on g5Rp-InsP_6_ interaction, amino acid residues involved in InsP_6_ binding pockets were substituted by single point mutation (Q6A, K8A, K94A, K98A, K133A). Each mutant was tested their binding affinity of InsP_6_ by MST, respectively. Fig. 3*D*, and Fig. S6 showed that mutants resulted in a notable decrease in g5Rp-InsP_6_ interaction, what's more, the quintuple mutant Q6A/K8A/K94A/K98A/K133A totally lost the binding ability with InsP_6_. Taken together, the mutagenesis work indicates that positively charged residues Lys8, Lys94, Lys98, Lys133 forming a cluster to mediate the g5Rp-InsP_6_ interaction.

### Analysis of residues involved in g5Rp-RNA interfaces

To characterize RNA binding surface on g5Rp, we analyzed the electrostatic potential at the surface of the g5Rp, which indicated that three highly positively charged areas (Area I-III) may play roles in g5Rp-RNA interaction (Fig. S2A). Area Ⅰ is located on the helical domain, containing residues Lys94, Lys95, Lys98, Arg100, Lys101 located on helix α5. Area Ⅱ compose of residues Lys8, Lys131, Lys133, Lys135, Arg146, Lys175, Lys179, and His180 mostly located on the β-strand 1,3, which close to the Nudix motif; Area III is located the very end of C-termius of g5Rp, comprising residues Arg^221^, Lys225, Arg^226^, Lys^243^, and Lys^247^ on helices α10 and α11 (Fig. 4*A*). To identify the mRNA binding surfaces on the g5Rp further, the residues mentioned above located in three positively charged areas of g5Rp were mutated respectively. The EMSA pattern showed that some mutants reduce the RNA binding affinity of g5Rp. Specifically, residues Lys^8^, Lys ^94^, Lys ^95^, Lys ^98^, Lys ^131^, Lys ^133^, Lys ^175^, Arg^221^, and Lys ^243^ were critical for single stranded RNA binding, albeit with different efficiencies (Fig. 4*B* and *C*), implying that the g5Rp–RNA interaction interfaces are mainly located at area Ⅰ, Ⅱ, and Ⅲ. These results also agree with our hypothesis that residues Lys^8^, Lys^94^, Lys^98^, Lys^133^ of g5Rp involved in both RNA and InsP_6_ interaction.

**FIG 4.**
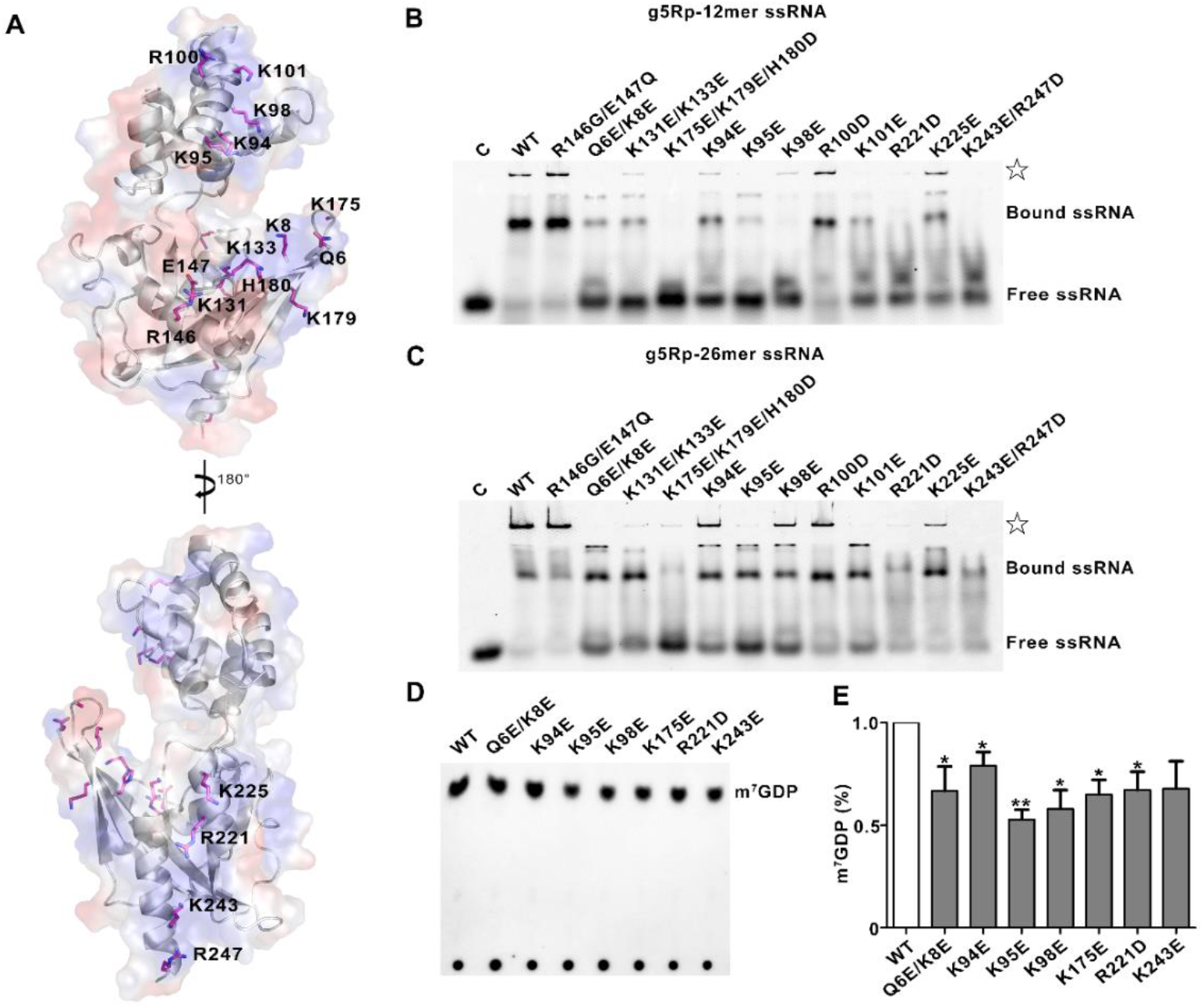
The residues involved in g5Rp-mRNA interfaces and decapping activity. **(A)**The mutation sites are located in the positive charged regions of g5Rp. **(B-C)** The binding abilities of g5Rp mutants to 12-mer and 26-mer ssRNAwere determined by EMSA, in the reaction, the protein concentration is 2.00u M, and the nucleic acid concentration is 0.25 uM, ☆ represents the precipitation in the gel **(D)** The decapping activity of g5Rp mutants. (E) The Semi-quantitative of m^7^GDP by Image J and drawing by graphpad prism8. (mean ± SD, n ≥ 3, *P < 0.05, **P < 0.01, unpaired t test).

We further explore that these key residues were responsible for cap cleavage in a manner dependent on the RNA moiety interaction. Mutant proteins including Q6E/K8E, K94E, K95E, K98E, K175E, R221D, and K243E were expressed and purified. Consistent with our previous data, incubation of the ^32^P-cap-labeled RNA substrate with wild-type g5Rp resulted in cap cleavage, as observed by m^7^GDP release. When equivalent amounts of the mutant proteins of the g5Rp were included in the decapping reaction, the amount of m^7^GDP released is reduced variously in each lane. Mutant K95E decrease the decapping activity almost 50% (Fig.4*D* and *E*), indicating that these residues of g5Rp play pivotal role in mRNA decapping by interacting with substrates mRNA.

### Residues Gly^132^, Lys^133^, and Glu^147^ in Nudix motif impact the decapping activity

The Nudix motif of hydrolases contains crucial residues involved in catalytic activity. However, the residues in the catalytic pocket of g5Rp are still elusive from view of structure. To elucidate the function of the key residues in g5Rp, three substrate-binding structures from the Nudix superfamily were selected to identify homologous domains with high similarity at the potential catalytic pockets (Fig. S7*A*, *B* and *C*), as shown in Supplementary Table 2 (viz., Ap4A hydrolase (*Aquifer aeolicus*, PDB: 3I7V) ^33^, nudix hydrolase DR1025 (*Deinococcus radiodurans*, PDB: 1SZ3), and MTH1 (*Mus musculus*, PDB: 5MZE) ^34,35^, (all belonging to the Nudix superfamily). Superposition of the C-terminus of g5Rp with that of MTH1, Ap4A hydrolase, and Nudix hydrolase DR1025 resulted in Cα backbone root-mean-square deviation values of 0.50, 3.08, and 5.6 Å, respectively, despite the low sequence identities among these proteins (Fig. S7*D*). Therefore, the potential substrate-binding site of g5Rp was proposed on the basis of the superpositions of these substrate-binding protein structures in the Nudix superfamily. Residues Gly^132^, Lys^133^, and Glu^147^ located on the Nudix motif of g5Rp may be responsible for cap cleavage.

To investigate the potential roles of these key residues located on the Nudix motif in the decapping activity, we substituted g5Rp residues G132, K133 and E147 from Nudix motif (Figure. 5*A* and S7*D*) with Ala, Glu, and Gln, respectively. As expected, the substitution of the residue K133 by glutamate resulted in a 30% decrease in the decapping activity. And the substitution of the residues G132, E147 by alanine and glutamine respectively, inactivate the decapping function of g5Rp completely (Fig.5*B* and *C*). No m^7^GDP was observed when the two mutants of the g5Rp were included in the decapping reaction, validating that the decapping activity was dependent on these two key residues located in Nudix hydrolase motif. Interestingly, EMSA results showed that mutant K133E reduces g5Rp's binding affinity to RNA which suggests that the loop region of Nudix motif takes part in substrates mRNA binding. (Fig.5*D* and *E*)

**FIG 5.**
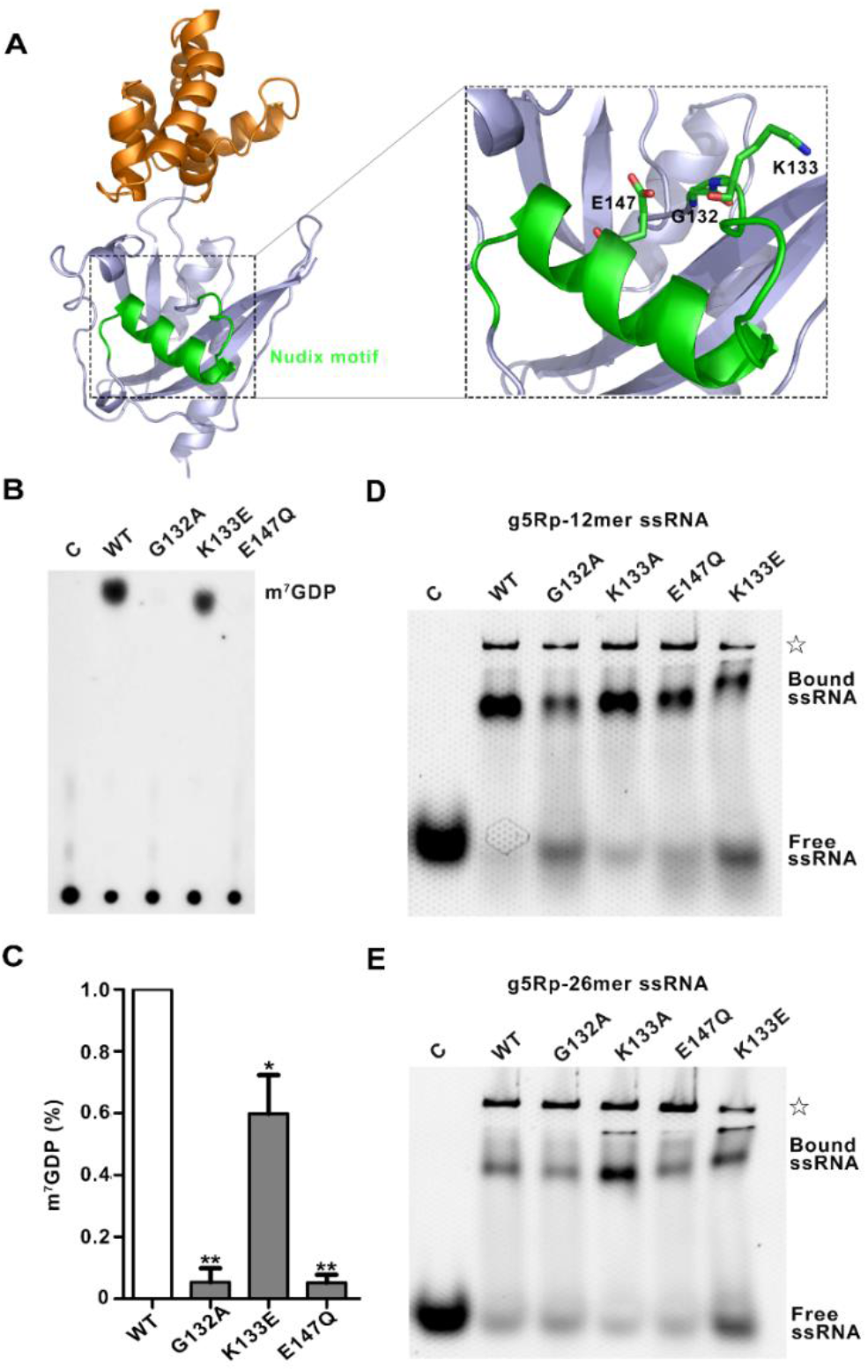
The Nudix motif of g5Rp involved in mRNA binding and decapping. (**A**)The details of nudix motif in g5Rp. The Nudix motif is colored by green. The mutant residues are shown as sticks. (**B**) The decapping activity of g5Rp mutants. (**C**) The Semi-quantitative of m^7^GDP by Image J and drawing by GraphPad prism8. (mean ± SD, n ≥ 3, *P < 0.05, ***P < 0.001, unpaired t test). (**D-E**) The binding abilities of g5Rp mutants to 12-mer and 26-mer ssRNsA were determined by EMSA,in this reaction, the protein experimental concentration is 2.00uM, and the nucleic acid experimental concentration is 0.25uM, ☆ represents the precipitation in the gel.

### InsP_6_ inhibits the decapping activity by disrupting g5Rp-mRNA interaction

The finding that residues located on mRNA binding regions of g5Rp are also playing pivotal roles in g5Rp-InsP_6_ interaction suggests that the InsP_6_ may inhibit the g5Rp decapping activity through preventing g5Rp from binding to its mRNA substrate (Fig. 6*A*). This prediction was confirmed by decapping and EMSA assays using recombinant g5Rp protein, InsP_6_ and RNA substrates *in vitro*. The increasing amounts of InsP_6_ were added to the decapping reactions to analyze its effect on RNA decapping by g5Rp. As shown in Fig. 6*B*, the addition of InsP_6_ significantly affected g5Rp cleavage, suggesting that the InsP_6_ can inhibit the decapping activity of g5Rp *in vitro*. To investigate if this inhibitory mechanism of InsP_6_ on g5Rp is due to inositol phosphate competitively inhibiting mRNA binding to the g5Rp, we further measured the competition of InsP_6_ with nucleic acids for the binding to g5Rp by using EMSA. As expected, the amount of free single-stranded nucleic acids accumulated with an increasing concentration of InsP_6_, demonstrating that InsP_6_ interrupts the g5Rp-mRNA interaction through the directly binding to g5Rp (Fig. 6*C* and *D*). In addition, all residues involved in InsP_6_ interaction in g5Rp were mutated to alanine at the same time. The quintuple mutant (Q6A/K8A/K94A/K98A/K133A) of g5Rp lost most of its ability to bind with both InsP_6_ and RNA, (Fig. S6*F* and S8*A*). These mutations also significantly affected the catalytic ability of g5Rp *in vitro* (Fig. S8*C* and *D*), suggesting that InsP_6_ inhibits the mRNA decapping activity of g5Rp through competing for the substrate mRNA binding surface in g5Rp.

**FIG 6.**
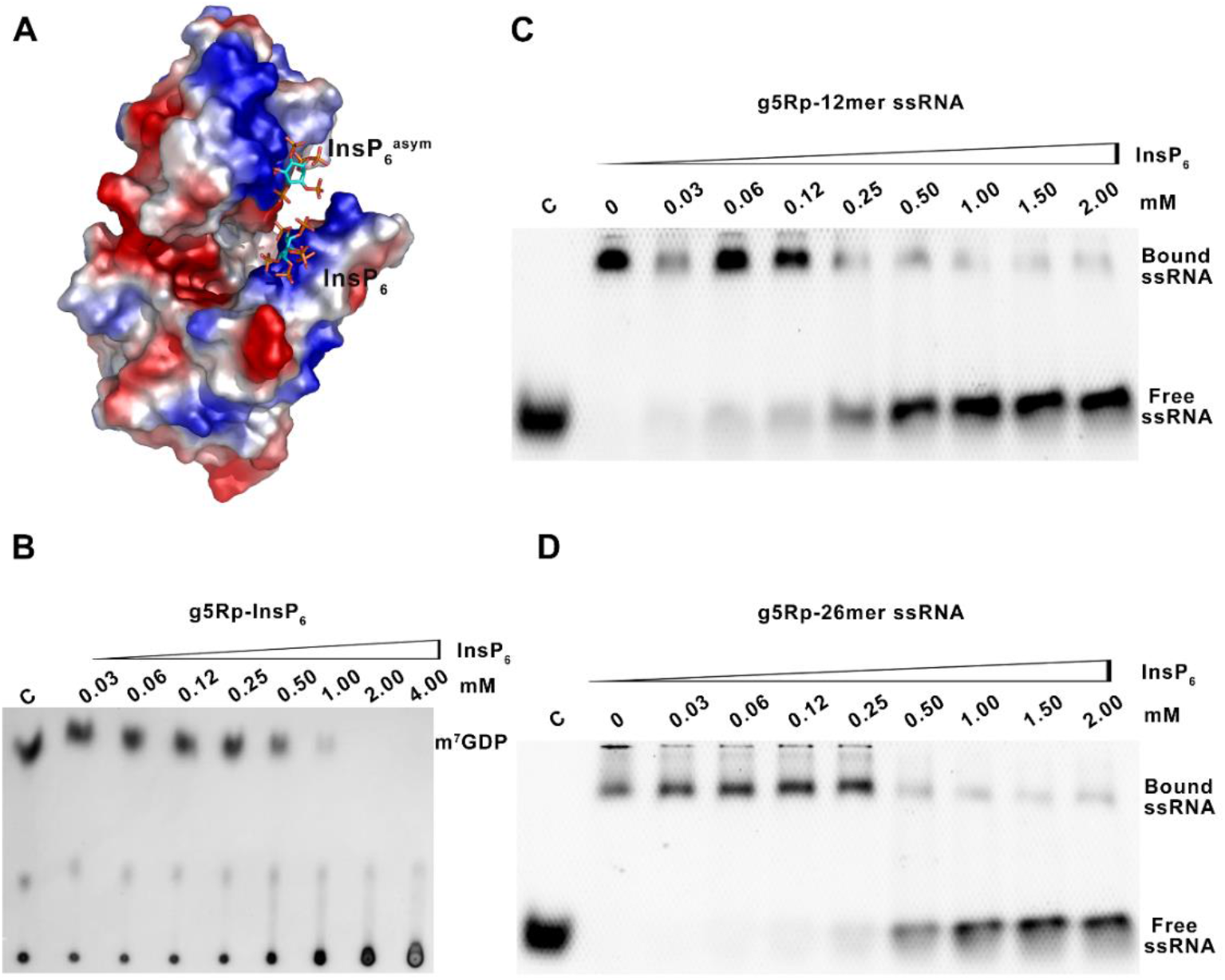
The influence of InsP_6_ on mRNA binding and decapping activity of g5Rp. **(A)** The surface charge of g5Rp in complex with InsP_6_. **(B)** The influence of InsP_6_ on decapping activity of g5Rp. (**C-D**) The influence of InsP_6_ on binding abilities of g5Rp with 12-mer and 26-mer ssRNA were determined by EMSA, respectively. The different concentrations of InsP_6_ with g5Rp in the reaction system is indicated above of the gel, the concentration of protein and the nucleic acid is 2.00uM and 0.25uM, respectively.

### Transient expression g5Rp decreases levels of mRNA substrates in 293T cells

The above data provide strong *in vitro* evidence for g5Rp-mRNA interaction being a critical step for the decapping enzyme process. To determine whether changes in g5Rp-mRNA interaction was directly related to the stability of cellular mRNAs *in vivo*, representative cellular mRNA (eIF4E, eIF4EA, TP53) levels were tested by RT-qPCR in cells. In 293T cells, the Flag-tagged g5Rp, and the g5Rp mutants (K8E, K94E, K95E, K98E, G132A, K133E, E147Q, K175E, R221D, and K243E) were overexpressed, respectively. As shown in Figure 7*A*, the proteins of g5Rp-WT and mutants were detected by western blot. The mRNA levels of target genes (eIF4E, eIF4EA, TP53) were decreased in 293T cells overexpressing g5Rp-WT. There are not obviously mRNA levels changed in catalytic destructive mutants Q132A and E147Q. The cells overexpressed truncated version g5Rp-ΔN and mutant K95E, R221D, which mutants significant lost the RNA binding ability *in vitro*, have no effect on the mRNA levels of target genes in 293T cells. Mutants K8E, K133E which declined the RNA binding *in vitro* have varying degrees of ascent compared with the mRNA levels of g5Rp-WT group. However, the changes in mRNA level of targets genes observed in mutants K94E, K98E, K175E and K243E have not statistical difference with g5Rp-WT (Fig. 7*B-D*). Taken together, these results suggest that key residues K8, K95, K133, and R221 playing pivotal roles in g5Rp-RNA interaction are also important to the g5Rp related cellular RNA degradation *in vivo*.

**FIG 7.**
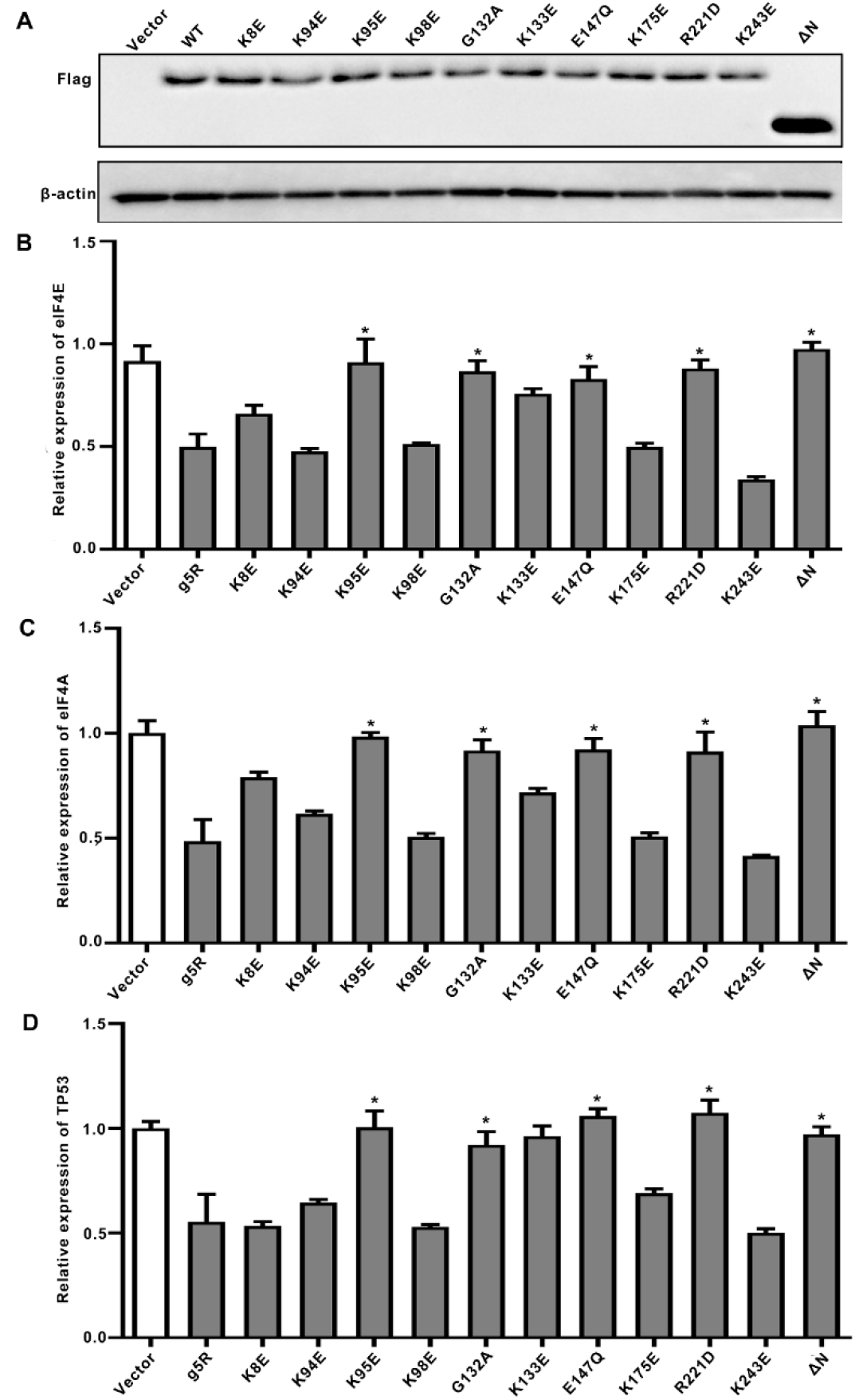
The influence of g5Rp and mutants on mRNA substrates level in 293T cells. **(A)** The expression of various g5Rp mutants in 293T cells were analyzed by western blot. **(B)** The mRNA levels of eIF4E in 293T cells when a series of g5Rp mutants overexpressed. **(C)**The mRNA level of eIF4A in 293T cells when a series of g5Rp mutants overexpressed. The mRNA level of TP53 in 293T cells when a series of g5Rp mutants overexpressed. (mean ± SD, n ≥ 3, *P < 0.05, **P < 0.01, unpaired t test).

### Discussion

Given that an ASFV outbreak in China would potentially devastate the world's largest pork producer, significant efforts have been made to determine the structures and functions of essential viral proteins that may be used as targets for new anti-ASFV drugs. Several structures of ASFV-encoded enzymes and associated proteins that are involved in viral transcription and replication have been reported, including AP endonuclease ^36^, the histone-like protein pA104R ^37^, pS273R protease ^38^, DNA ligase ^39^, and dUTPase ^40,41^. However, the structures and functions of some critical ASFV proteins remain elusive, including those of g5Rp, a decapping enzyme that is crucial for viral infection ^23^. Our structures of g5Rp alone and in complex with InsP_6_ provide the molecular basis for g5Rp substrate recognition and reveal that inositol phosphate was involved in the regulation of cellular mRNA degradation through direct interaction with the ASFV decapping enzyme g5Rp. Three potential RNA binding regions are identified, including a novel folding domain located on the N-terminal of g5Rp and Nudix motif on its C-terminus. More importantly, identification of the major nucleic acid-binding surfaces as well as the binding pocket of InsP_6_ on g5Rp provides important structural information and a novel strategy for future anti-ASFV drug design.

To explore the nucleic acid binding properties of g5Rp, we conducted a series of nucleic acid binding experiments. Results indicating that an intact dimeric interface is efficient for g5Rp-RNA interaction. Meanwhile, both N-termius (helical domain) and C-terminal (Nudix domain) of g5Rp are involved in ssRNA interaction. Our EMSA and SPR measurements show that the N-terminal domain of g5Rp can bind with ssRNA with equally high affinity compared with full-length. It composes of six α-helices forming a globin-fold-like domain, which is different from the traditional RNA-binding domain that prefers to adopt the alpha/beta topologies^42–45^. According to the g5Rp structure, the surface electrostatic potential characteristics of the N-terminus present a highly positive charged area on the helix α5. The single point mutations of positively charged residues in N terminus significantly reduced the nucleic acid-binding activity of g5Rp with ssRNA (Fig. 4*B* and *C*). Furthermore, there are two positive areas located on the C-terminus of g5Rp, including the Nudix motif, participating in the substrate RNA interaction. We mutated the two positively charged regions (K8A/K131A/K133A/K135A and R221A/K225A/R226A/K243A/R247A) located in the Nudix domain, the EMSA data showed that the nucleic acid binding ability of these two mutants was significantly reduced (Fig. S8*B*), and the figure S8*E* and S8*F* data showed that the substantially decline in capacity of K8A/K131A/K133A/K135A remove the m^7^GpppRNA cap. These results predicated that the Nudix motif of g5Rp possesses substrate-selectivity at the step of mRNA binding.

Previously studies revealed that the Nudix motif (residues 132–154) is an essential component of the α–β–α sandwich in catalytic center of g5Rp. Several of the conserved catalytic amino acids and glutamate residues (E^147^, E^149^, E^150^, and E^151^) located on the α-helix of the Nudix motif of g5Rp have been found to be important for the activity of Nudix hydrolases^23,28^. However, the function of the loop region within the Nudix motif is exclusive leading us to predicated that the loop region may contribute to binding with mRNA. Therefore, we mutated several residues in this loop region, including the mutations K133E, G132A, and examined the effects on the protein's interaction with single-stranded nucleic acids. It is interesting to find that substituted the conserved residues K^133^, G^132^ are highly sensitivity to g5Rp-RNA interaction. Comparing with glutamate residues located on the α-helix of the Nudix motif of g5Rp involved in mRNA cap structure interaction, residues K^133^, G^132^ are important to bind with RNA moiety on the substrate. In this way, we provided the demonstration that the short loop in the Nudix motif is required for g5Rp–RNA interaction. Including the Nudix motif, three positively charged patches on the g5Rp surface were mapped as mRNA-binding regions. Furthermore, we also investigated the importance of the mRNA–interacting residues in g5Rp mediated decapping. The g5Rp mutants K8E, K94E, K95E, K98E, K175E and R221D, showed a strong reduction in decapping activity, demonstrating the importance of the mRNA binding residues for catalysis. Therefore, further work should be done to elucidate the structure of g5Rp in complex with mRNA.

The other important finding in this study was that InsP_6_ is able to inhibit the decapping activity of g5Rp. As we know, InsP_6_ is widespread in cells with diverse biological functions^46–49^. Here, we found that InsP_6_ competes with mRNA substrates for binding to g5Rp and inhibits its decapping activity. Previously study reported that the g5Rp is a diphosphoinositol polyphosphate phosphohydrolase (DIPP), which preferentially remove the 5-β-phosphate from the InsP_7_ to produce InsP_6_ with unclear functional significance^25^. Later, Susan P. and colleagues identified that g5Rp can hydrolyze the mRNA cap when tethered to an RNA moiety *in vitro* ^23^. Our results show InsP_6_ the product of g5Rp playing the role of DIPP can directly inhibit the mRNA decapping activity of g5Rp. To illustrate the structural basis of the inhibitory mechanism of InsP_6_ to the decapping activity of g5Rp, we solved the complex structure of g5Rp with InsP_6_, also the enzyme-product complex in Nudix superfamily. To our surprised, the InsP_6_ is located on the mRNA binding region instead in the catalytic center of the g5Rp. Furthermore, we superposed the catalytic domain of g5Rp-InsP_6_ complex with the structures of human DIPP1 in complex with the substrate InsP_7_ ^50,51^. The visualizing result showed that the substrate InsP_7_ is located in the catalytic center of DIPP1 unlike the InsP_6_ sitting on the edge of catalytic domain of g5Rp (Fig S9*A*). Therefore, the structure of g5Rp-InsP_6_ complex may represent an intermediate in the release of the product of the enzymatic reaction^52^. This also explain why InsP_6_ binding decrease the temperature value (B factor) in the InsP_6_ binding sites compared with B factor in the same regions of the g5Rp wild-type structure (Fig. S9*B*). We also noticed that the InsP_6_ was refined with correspondingly high B-factor that exceed the average B-factor of protein in complex. Considering the g5Rp-InsP_6_ interaction with a dissociation constant Kd in the 22.5uM range, the ligand only achieves a reasonable occupancy of 70%^53^. To avoid instance of overenthusiastic interpretation of ligand density, we test the InsP_6_ binding site by using single point mutations. Residues involved in the InsP_6_ binding surface of g5Rp substituted by alanine (Q6A/K8A/K94A/K98A/K133A) reduced its InsP_6_-binding capacity and RNA interaction, indicating destructive InsP_6_ binding site has the capability to abolish the substrate RNA binding ability of g5Rp (Fig. S8*A*, *C* and *D*).

Our study raises the possibility that g5Rp hydrolyzes InsP_7_ to up-regulation of the level of InsP_6_, which is a key regulator of g5Rp-mediated mRNA decapping during the ASFV infection *in vivo*. Very recently, Soumyadip S. and colleagues reported that InsP_7_ regulates the NUDT3-medicated mRNA decapping and also observed the phenomenon that InsP_6_ inhibits mRNA decapping by NUDT3^54^. There are emerging signs that the functions of InsP_6_ are associated with mRNA transportation and degradation in ASFV infected cells. Further studies into the function of the InsP_6_ and the regulation mechanism in the inositol-based cell signaling family during the viral infection is required.

## MATERIALS AND METHODS

### Cell culture

The human 293T cells were cultured in Dulbecco's modified Eagle's medium (DMEM) (Hyclone) supplemented with 10% fetal bovine serum (FBS) (Gibco), 100 U/mL penicillin and 100 micro g/mL streptomycin at 37 °C under a humidified 5% CO2 atmosphere (Thermo).

### Plasmid construction, protein expression, and purification

The gene encoding ASFV g5Rp (*D250R*) was synthesized and subcloned into the pSMART-1 and pcDNA3.1 respectively. The point mutants and truncation variants of g5Rp (viz., Q6E/K8E, K94E, K95E, K98E, R100D, K101E, K131E/K133E, R146G/E147Q, K175E/K179E/H180D, R221D, K225E, K243E/R247D, Q6A/K8A/K94A/K98A/K133A, K8A/K131A/K133A/K135A, R221A/K225A/R226A/K243A/R247A, I84A/I116A/L200A/I206A/F222A, g5Rp-Δ C (Helical domain, residues 36-124) and g5Rp-Δ N (Nudix domain, connecting residues 1-35 and 125-250 directly)) were generated using the Fast Mutagenesis V2 Kit (Vazyme Biotech, China). The primers used in this study are listed in Supplementary Table 3. The recombinant plasmids were confirmed by sequencing (Sangon Biotech, China) before being introduced into *E. coli* BL21 (DE3) (Invitrogen, US) or human 293T cells. The bacterial cells were cultured in Luria Broth medium at 35°C until the optical density at 600 nm reached 0.6–0.8. Protein expression was then induced by the addition of isopropyl β-D-1-thiogalactopyranoside for 16 h at 16°C. The g5Rp molecules were purified by Ni-NTA (Qiagen, Germany) affinity chromatography, followed by heparin affinity chromatography (GE Healthcare, US). The peak fractions containing the target proteins were pooled, concentrated to 1 mL, and finally loaded onto a Superdex 75 column (GE Healthcare, US) for further purification and characterization. Selenomethionine-labeled g5Rp (SeMet-g5Rp) was then prepared using a previously described protocol ^55^. The purity of all proteins was above 95% on the SDS-PAGE gel.

### Protein crystallization and optimization

The prepared SeMet-g5Rp was concentrated to 12 mg/mL for the crystallization trials. The crystals were grown using the hanging-drop vapor diffusion method at 16°C in a reservoir solution containing 0.1 M sodium citrate tribasic dihydrate (pH 5.8), 0.54 M magnesium formate dihydrate, and 10% (v/v) 1,2-butanediol as an additive reagent. The g5Rp–InsP_6_ complexes were prepared by mixing g5Rp with InsP_6_ at a stoichiometric ratio of 1:3. Then, using the hanging-drop vapor diffusion method, crystals of the complexes were grown from 1 M imidazole (pH 7.0) at 16°C. All crystals were transferred into solutions containing 20% (v/v) glycerol prior to being frozen and stored in liquid nitrogen.

### Data collection, processing, and structure determination

The single wavelength anomalous dispersion (SAD) data were collected using synchrotron radiation of 0.98 Å wavelength under cryogenic conditions (100K) at BL18U1 beamline, Shanghai Synchrotron Radiation Facility. All diffraction data sets including g5Rp-WT and complex with InsP_6_ were indexed, integrated and scaled by using HKL-2000 package ^56^. The selenium atoms in the asymmetric unit of SeMet-g5Rp were located and refined, and the SAD data phases were calculated and substantially improved through solvent flattening with the PHENIX program ^57^. A model was built manually into the modified experimental electron density using the model-building tool Coot ^58^ and then further refined in PHENIX. The model geometry was verified using the program MolProbity ^59^. Molecular replacement was used to solve the structure of the g5Rp–InsP_6_ complex, using Phaser in the CCP4 program suite with an initial search model of SeMet-g5Rp ^60^. Structural figures were drawn using PyMOL (DeLano Scientific). The data collection and refinement statistics are shown in Table 1.

**TABLE 1.**
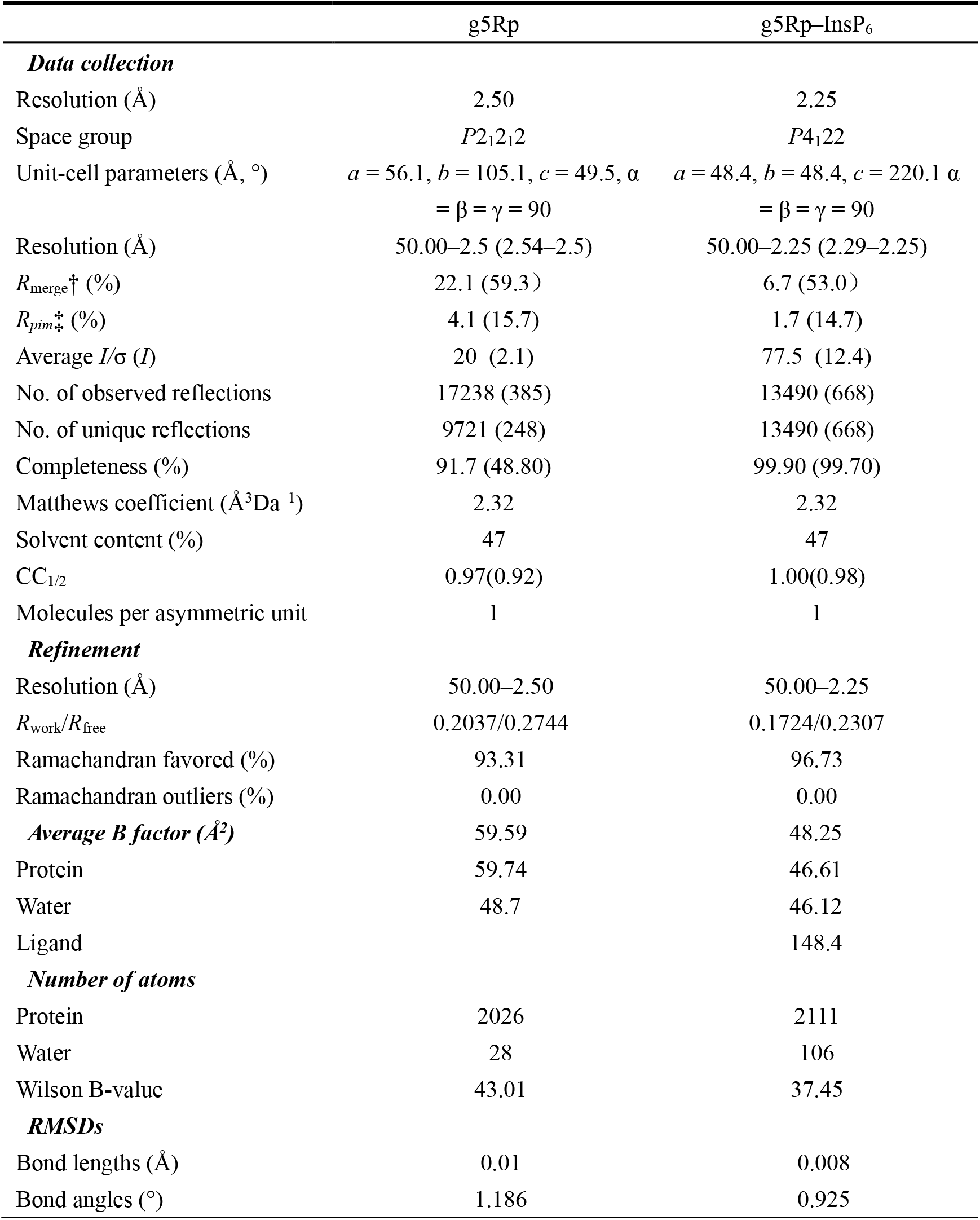
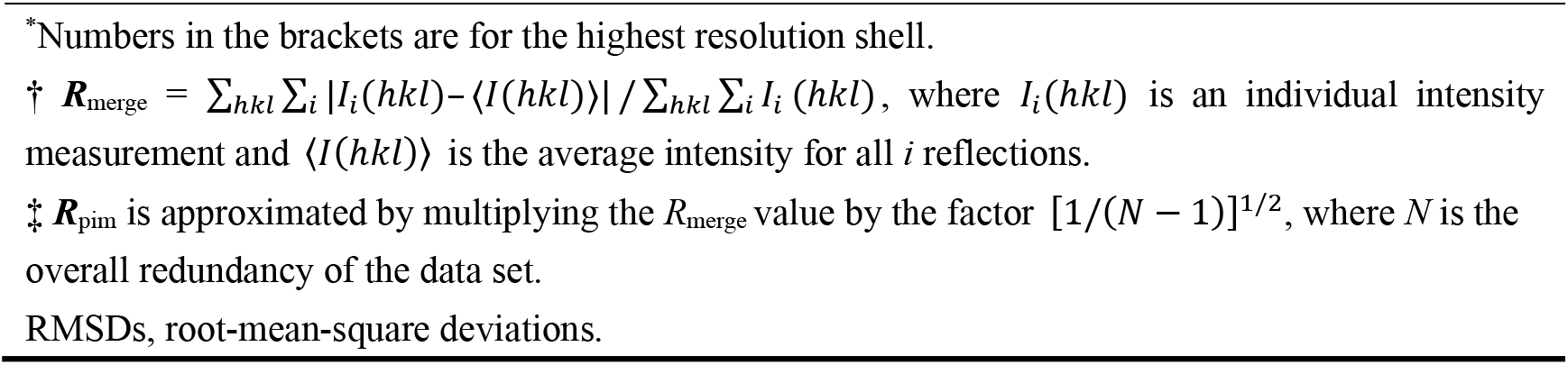
Data collection and refinement statistics

|  | g5Rp | g5Rp–InsP<sub>6</sub> |
| --- | --- | --- |
| **Data collection** |  |  |
| Resolution (Å) | 2.50 | 2.25 |
| Space group | $P2_12_12$ | $P4_122$ |
| Unit-cell parameters (Å, °) | $a = 56.1, b = 105.1, c = 49.5, \alpha = \beta = \gamma = 90$ | $a = 48.4, b = 48.4, c = 220.1, \alpha = \beta = \gamma = 90$ |
| Resolution (Å) | 50.00–2.5 (2.54–2.5) | 50.00–2.25 (2.29–2.25) |
| $R_{\text{merge}}^{\dagger}$ (%) | 22.1 (59.3) | 6.7 (53.0) |
| $R_{\text{pim}}^{\ddagger}$ (%) | 4.1 (15.7) | 1.7 (14.7) |
| Average $I/\sigma$ ($I$) | 20 (2.1) | 77.5 (12.4) |
| No. of observed reflections | 17238 (385) | 13490 (668) |
| No. of unique reflections | 9721 (248) | 13490 (668) |
| Completeness (%) | 91.7 (48.80) | 99.90 (99.70) |
| Matthews coefficient (Å<sup>3</sup>Da<sup>−1</sup>) | 2.32 | 2.32 |
| Solvent content (%) | 47 | 47 |
| CC<sub>1/2</sub> | 0.97(0.92) | 1.00(0.98) |
| Molecules per asymmetric unit | 1 | 1 |
| **Refinement** |  |  |
| Resolution (Å) | 50.00–2.50 | 50.00–2.25 |
| $R_{\text{work}}/R_{\text{free}}$ | 0.2037/0.2744 | 0.1724/0.2307 |
| Ramachandran favored (%) | 93.31 | 96.73 |
| Ramachandran outliers (%) | 0.00 | 0.00 |
| **Average B factor (Å<sup>2</sup>)** | 59.59 | 48.25 |
| Protein | 59.74 | 46.61 |
| Water | 48.7 | 46.12 |
| Ligand |  | 148.4 |
| **Number of atoms** |  |  |
| Protein | 2026 | 2111 |
| Water | 28 | 106 |
| Wilson B-value | 43.01 | 37.45 |
| **RMSDs** |  |  |
| Bond lengths (Å) | 0.01 | 0.008 |
| Bond angles (°) | 1.186 | 0.925 |

\*Numbers in the brackets are for the highest resolution shell.
† $R_{\text{merge}} = \sum_{hkl} \sum_i |I_i(hkl) - \langle I(hkl) \rangle| / \sum_{hkl} \sum_i I_i(hkl)$ , where $I_i(hkl)$ is an individual intensity measurement and $\langle I(hkl) \rangle$ is the average intensity for all $i$ reflections.
‡ $R_{\text{pim}}$ is approximated by multiplying the $R_{\text{merge}}$ value by the factor $[1/(N - 1)]^{1/2}$ , where $N$ is the overall redundancy of the data set.
RMSDs, root-mean-square deviations.

### Surface plasmon resonance analysis

The SPR analyses were carried out using the BIAcore 8K system with a streptavidin-coated (SA) chip (#BR-1005-30, GE Healthcare) at 16°C. To reduce effects attributed to mass transport, low levels of biotin-labeled ssRNA (−5′ GCUUUGAUUUCGUGCAUCUAUGGAGC-3′), (5′-GCUUUGAUUUCG-3′) ligands (given in relative unites [RU]) were immobilized on the SA chip. Given the apparent variation in RU of immobilized ligand used in different binding studies^61,62^, the average RU of immobilized 26mer RNA and 12mer RNA in this study are approximately 200-400RU. The blank channel served as a negative control. Protein solutions with varies concentrations run across the chip at a rate of 30 μL/min, then dissociated by running buffer (20 mM HEPES, 150 mM NaCl, 3mM EDTA and 0.05% Tween 20 (pH 7.5)) for 300 s at a flow rate of 30 μl/min. Regeneration of the sensor chips was performed for 30 s using regeneration buffer (0.5%SDS). The data on the binding of the g5Rp molecules to ssRNAs were fitted to a kinematic binding model, which determines association and dissociation constants by fitting the experimental data to a Langmuir model with 1:1 interaction model between analyte A and ligand B: the association (ka) and dissociation (kd) rate constants, and the affinity value 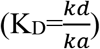 were determined.

### Microscale thermophoresis

The binding between g5Rp and InsP_6_ was measured by microscale thermophoresis. In brief, g5Rp was first labeled using the Monolith NT Protein Labeling Kit RED-NHS (NanoTemper Technologies), and the labeled protein was then diluted to 20 nM with buffer containing 50 mM HEPES, 300 mM NaCl, and 0.05% (v/v) Tween 20 (pH 7.0). Then, a series of concentrations of InsP_6_ diluted in a buffer composed of 50 mM HEPES, 300 mM NaCl, and 0.05% (v/v) Tween 20 (pH 7.0) was added. The mixtures were loaded into capillaries and measured at ambient temperature by using 20% LED and medium MST power in a Monolith NT.115 system (NanoTemper Technologies). The data were analyzed using NanoTemper analysis software (v.1.2.101).

### Electrophoretic mobility shift assay

EMSAs were performed to determine the nucleic acid affinity of the wild-type and g5Rp mutants. The single-stranded nucleic acids used were 6‐carboxyfluorescein‐labeled ssRNA(5′-GCUUUGAUUUCGUGCAUCUAUGGAGC-3′), 6‐carboxyfluorescein‐labeled ssRNA(5′-GCUUUGAUUUCG-3′) (Sheng Gong, Shanghai, China). Initially, 0.25 μM of the ssRNA was incubated on ice for 30 min together with different concentrations of wild-type in a buffer composed of 20 mM HEPES, 60 mM KCl, 0.5 mM EDTA, 0.1% Triton-X-100, 4mM DTT, 2mM MgCl_2_ and 5% (v/v) glycerol (pH 7.9), the experimental concentration of mutants g5Rp is 2uM. To determine the effect of InsP_6_ on nucleic acid-binding ability of g5Rp, we also used the EMSA to test the nucleic acid-binding ability of the enzyme mixed with different concentrations of InsP_6_ (0.03mM, 0.06mM, 0.12mM, 0.25mM, 0.5mM, 1mM, 1.50mM, 2.00mM). All samples were incubated on ice for 30 min and then electrophoresed on 4.0 % native‐PAEG gels for 45 min at a voltage of 100 V. The results were determined with a Bio-Rad ChemiDoc™ MP Imaging System (Bio-rad, USA).

### Capping of the mRNA Body

For decapping assays with g5Rp, uncapped RNA was transcribed using the T7 RNA transcription kit (Vazyme) with the linear DNA(5’-CATTATTGGCCTGAAAAAGATGATTGACAGCTATAATGATTACTACAACAACGAAGTTTTCGTTAAACATAAAAACC-3’) as a template. The RNAs were capped in 50ul reaction system typically containing 0.4 nmol RNA, 12ul 3000Ci/ mmol [α-^32^P] GTP, 0.67 mM Sadenosylmethionine, 50 mM Tris–HCl pH 7.6, 2 mM MgCl_2_, 6mM KCl, 1 mM DTT and 75 U Vaccinia Capping Enzyme (NEB) at 37°C for 2 h. Cap-labeled RNAs were then separated from free[α-^32^P]GTP nucleotide by chromatography through a G-50 column(sigma).

### mRNA Decapping Assays

All decapping experiments were carried out in a buffer containing 50 mM Tris-HCl (pH 7.0), 1 mM DTT and 2 mM MnCl_2_. Generally, 100ng of wild-type or g5Rp mutants was used in each 5μl reaction. To determine the correlation between InsP_6_ and decapping ability of g5Rp, we also test the decapping ability of the enzyme mixed with different concentrations of InsP_6_ (0.03mM, 0.06mM, 0.12mM, 0.25mM, 0.5mM,1mM,2mM,4mM). The decapping reaction was carried out at 37°C for 60 min, then stopped by 25 mM EDTA. The products of the reaction were separated by PEI-cellulose thin layer chromatography developed in 0.45M (NH4)2SO4 and detected with autoradiography.

### Western Blotting and Analysis

200,000 cells/well were seeded in 6-well plate (Nest) and cultured for 24h. Then plasmids (2 μg/well) were transfected into cells using Lipofectamine® 3000 (Invitrogen). After 48 h cells were lysed in RIPA (Biosharp) containing protease inhibitor Cocktail (MCE) and quantified by the Bicinchoninic Acid (BCA) kit (Sangon). About 20 μg protein was loaded in 12% SDS-PAGE and transferred to PVDF (0.22 μM) membrane (Millipore). Membranes were incubated with primary antibodies including anti-β-actin (1:10,000 dilution; HuaBio) and anti-Flag (1:1,000 dilution; CST) at 4 °C overnight. After the membranes were washed three times for 10 min with TBST and incubated with secondary antibodies for 2 h at room temperature, they were washed three times again in TBST. Then the blots were detected with enhanced chemiluminescence reagents (Millipore) using MiniChemi™ (Sagecreation, China)

### RNA extraction and quantitative real-time PCR

Briefly, 100,000 cells were seeded in a 12-well plate (Nest) and cultured for 24h before transfection plasmid (1 μg/well). Total RNA was extracted using RNAiso Reagent (TaKaRa), and 500 ng of total RNA was reversely transcribed using PrimeScriptTM RT Reagent Kit (Takara). Subsequently, real-time polymerase chain reaction (PCR) amplification was performed with SYBR Premix ExTag (Takara) on a QuantStudio™ 3 System (Applied Biosystems). The cycling settings were as follows: 95 ℃ for 30 s, then 40 cycles of 95 ℃ for 5 s and 60 ℃ for 34 s. Relative mRNA levels were determined using the 2 -∆∆Ct (∆∆Ct =ΔCT (test)-ΔCT(calibrator)) method.

## Data availability

The structure factors and atomic coordinates of apo g5Rp and g5Rp-InsP6 have been deposited in the Protein Data Bank under the PDB ID codes 7DNT and 7DNU, separately.

## Acknowledgements

We thank the staff at the State Key Laboratory of Biotherapy, Sichuan University, who assisted with our research work during the period of the COVID-19 epidemic. The X-ray diffraction experiments were carried out at the Shanghai Synchrotron Radiation Facility (SSRF) at BL18U1. We also thank the beamline staff for their technical help during the data collection. We appreciate helpful advice and discussion with Dr. Xiaoyu Xue in Texas State University. This work was supported by grants from the National Key Research and Development Program of China (2017YFA0505903, 2018YFC1312300), the Special Research Fund on COVID-19 of Sichuan Province (2020YFS0010), and the Key Project on COVID-19 of West China Hospital, Sichuan University (HX-2019-nCoV-044).

