## Supplementary material for "Structural insight into molecular inhibitory mechanism of InsP_6_ on African Swine Fever Virus mRNA-decapping enzyme g5Rp": Supplementary information.pdf

### Supplementary results

#### Structures determination

The selenium-labeled wild-type g5Rp was expressed in *E. coli* cells and purified by nickel-nitrilotriacetic acid (Ni-NTA) (Qiagen) affinity chromatography following gel-filtration chromatography. The purified wild-type g5Rp, truncated g5Rp, and a series of g5Rp mutants were analyzed by circular dichroism (CD) spectroscopy to characterize the protein folding before any experiments (Fig. S10*A*-*C*). The selenium-labeled protein was screened for crystallization conditions and yielded crystals for X-ray diffraction analysis (Fig. S10*D*). The crystal structure of g5Rp was determined with the single-wavelength anomalous diffraction (SAD) method (1) at a resolution of 2.5 Å (Fig. S10*F*), with a final $R_{work}/R_{free}$ value of 0.20/0.27. The atomic model had a suitable geometry, with 97.5% of the residues located in the favored and allowed regions of the Ramachandran plot (2). The final g5Rp structure revealed one molecule in the asymmetric unit at space group $P2_12_12$. The structure of g5Rp is in the full-length polypeptide, except the residues Met1–Met5, for which the electron density map was ambiguous and assumed to be disordered in the crystals. The crystal shape of the g5Rp-$InsP_6$ complex is different from apo Se-g5Rp (Fig. S10*E*). The structure of g5Rp in complex with $InsP_6$ was determined using the molecular replacement method (3), where the initial model of g5Rp was solved by SAD and refined to 2.25 Å resolution (Fig. S10*G* )in space group $P4_122$ with a final $R_{work}/R_{free}$ value of 0.17/0.23. The Matthews coefficient for the g5Rp–$InsP_6$ complex revealed that there was only one protein complex in an asymmetric unit (4).

### Supplementary materials and methods

#### Cross-linking assay

A cross-linking assay was carried out by incubating 1 mg/mL of wild-type g5Rp or each g5Rp truncation variant and mutant I84A/I116A/L200A/I206A/F222A in a buffer containing 20 mM HEPES, 200 mM NaCl, 1 mM dithiothreitol, and 10% (v/v) glycerol (pH 7.5) and different concentrations of ethylene glycol *bis* (succinimidyl succinate) for 15 min at 4℃, EGS is dissolved in 100% DMSO, its reservoir concentration is 25mM. The reaction was terminated by adding glycine at a final concentration of 0.15 M. The reaction products were separated by 12% SDS-PAGE and detected by Coomassie Brilliant Blue staining.

#### Circular dichroism spectroscopy

Before CD measurements, diluting all purified proteins with purified water to 0.1mg/ml. The CD spectra were recorded on a Jasco J–715 Spectropolarimeter (JASCO, MD, United States) with the use of three scans on average within the 190–240 nm wavelength range from solutions in 2 mm path length quartz cuvettes at 25°C. Raw ellipticity data θobs (in 104 millidegrees) were converted to mean residue ellipticity (θ) (in 104 millidegrees) using the formula,

$$(\theta)=\frac{(\boldsymbol{\theta obs}\times \boldsymbol{MRW}\times 100)}{(\boldsymbol{c}\times \boldsymbol{l})}$$

where θ is the ellipticity in 104 millidegrees, l is the path length of the cuvette in cm, MRW is the mean residue weight, and c is the concentration in mg/ml. Data were analyzed with Microsoft Office Excel and Origin 8.

### Supplementary Figures

**Supplementary figure 1 The biochemical characters of the African swine fever virus decapping enzyme g5Rp.** (**A**) The elution profile of g5Rp in superdex200$^{TM}$ 10/300 column, the molecular weight of the standards and g5Rp are showed on this picture. (**B**) SDS-PAGE electrophoresis result of g5Rp. M is the protein maker. (**C**)The crosslinking of g5Rp by using chemical reagent EGS. The different concentrations of EGS with g5Rp in the reaction system is indicated above of the gel. (**D-E**) Superposition of g5Rp Nudix domain with DCP2 from S. pombe (mRNA decapping complex subunit 2, PDB: 5KQ4), DCP2 from K.lactis (Enhancer of mRNA-decapping protein 3, PDB: 6AM0), g5Rp ASFV is shown in brightorange, 5KQ4 is shown in palecyan and 6AM0 is shown in lightblue.

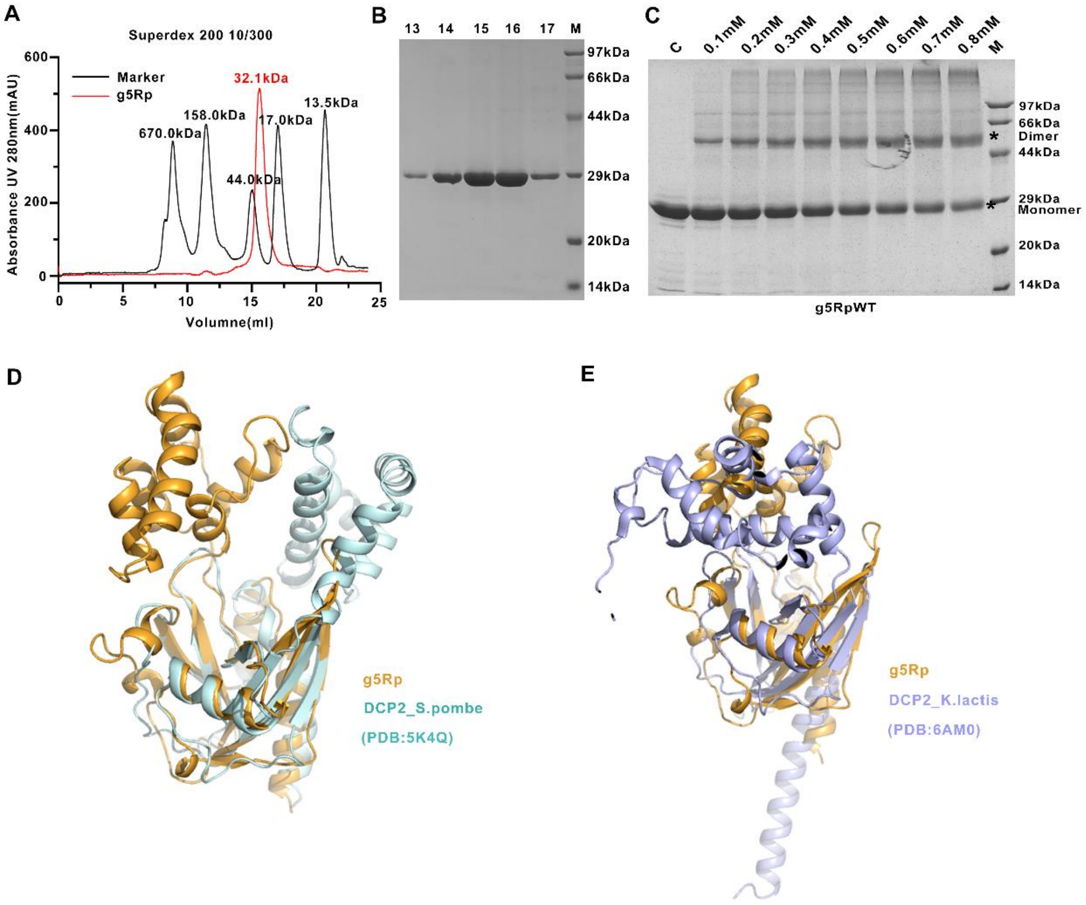

**Supplementary figure 2 Surface charge distribution of g5Rp and the Biochemical information of the African swine fever virus decapping enzyme truncated g5Rp**. (**A**) Surface charge distribution of g5Rp. The range of electrostatic surface potential is shown from -74.712kT/e in red color to +74.712 kT/e in blue color. The three highly positively charged areas are marked by the black dotted lines. (**B**) The binding abilities of g5Rp truncates (the final concentrations of WT and ΔC are 2.00μM while ΔN is 50uM) to 26- and 12-mer ssRNAs (final concentration 0.25 μM) were determined by EMSA. (**C** and **D**) The binding abilities of truncated g5RpΔC to 26- and 12-mer ssRNAs were determined by SPR.

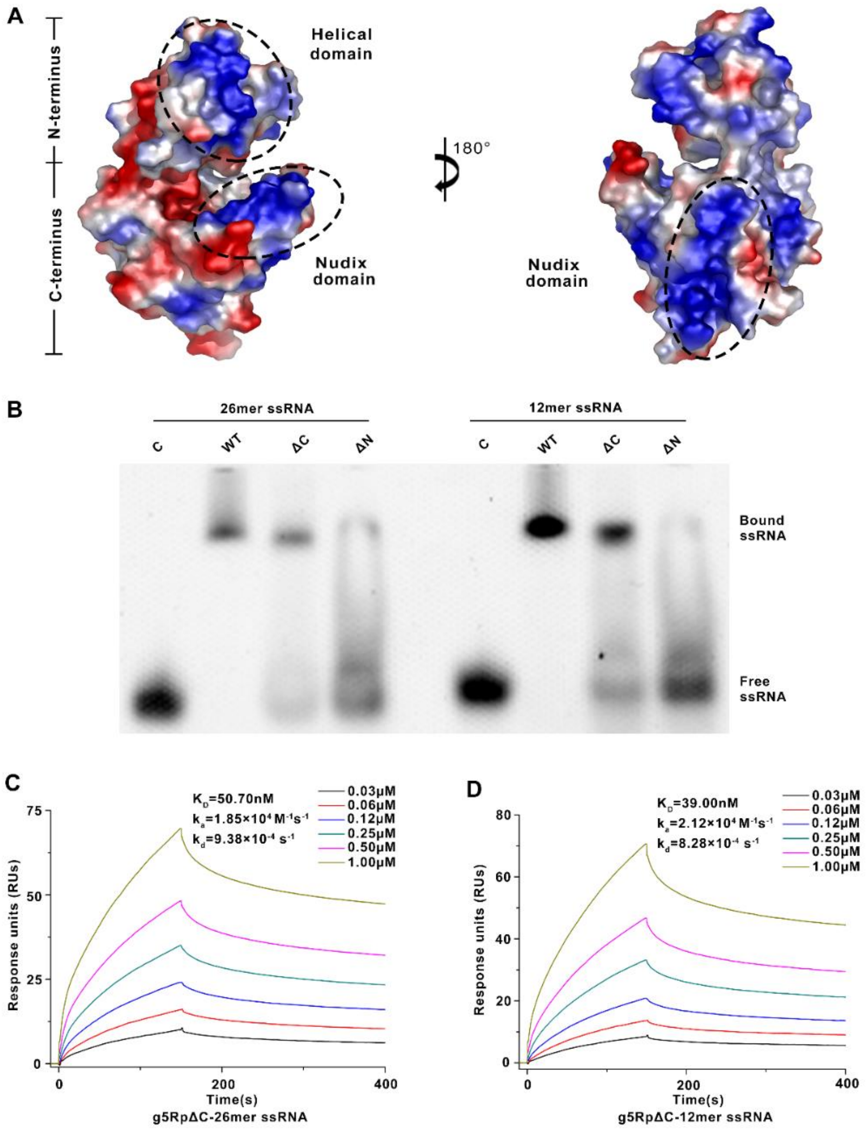

**Supplementary figure 3 The details structure information of dimeric g5Rp and the crosslinking result of truncated g5Rp.** (**A**) The details structure information of dimeric g5Rp. The residues of protomer A and B involved in interface are colored by purple and bule, respectively. (**B** and **C**) The crosslinking of truncated g5Rp by EGS. The different concentrations of EGS with g5Rp in the reaction system is indicated above of the gel.

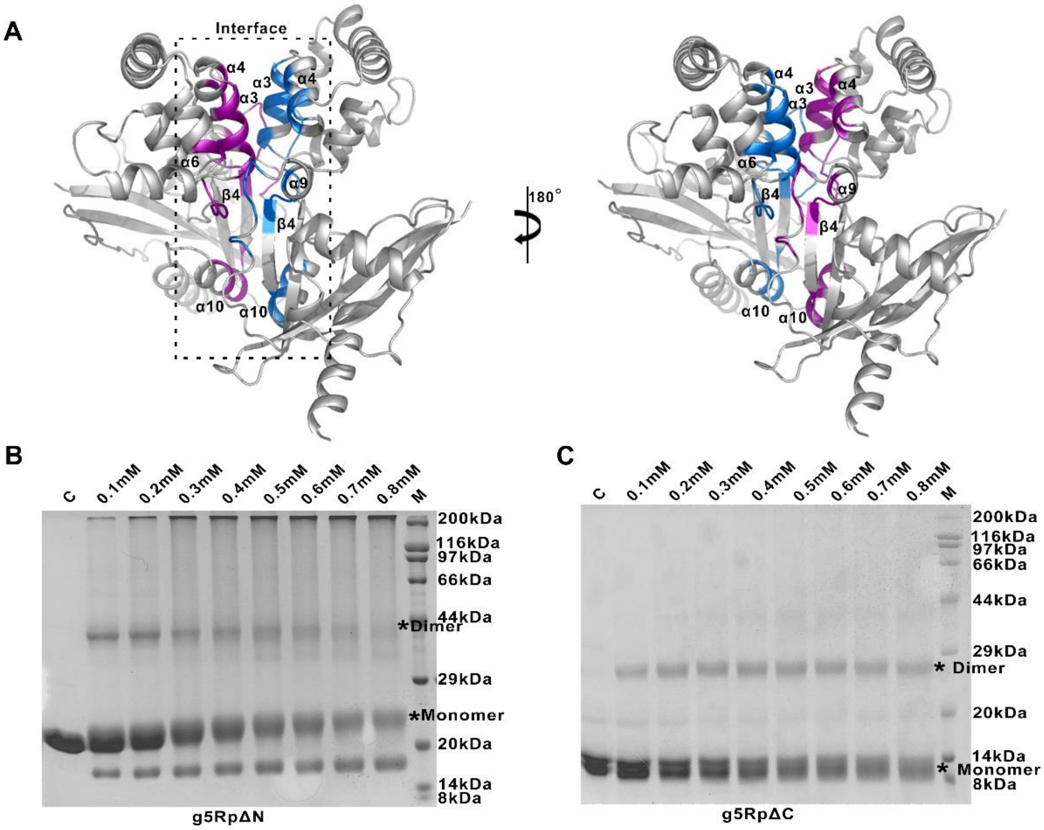

**Supplementary figure 4 The Biochemical information of the African swine fever virus decapping enzyme monomeric g5Rp.** (**A**) The elution profile of g5Rp-WT and g5Rp-I84A/I116A/L200A/I206A/F222A in superdex200$^{TM}$ 10/300 column, the molecular weight of the standards and g5Rp is showed on this picture. (**B**) The crosslinking of g5Rp-I84A/I116A/L200A/I206A/F222A by EGS. The different concentrations of EGS with mutant in the reaction system is indicated above of the gel. (**C**)The binding abilities of monomeric g5Rp (I84A/I116A/L200A/I206A/F222A) to 26- and 12-mer ssRNAs were determined by SPR. (**D**) The binding abilities of monomeric g5Rp (the final concentrations of WT and I84A/I116A/L200A/I206A/F222A are 2.00μM) to 26- and 12-mer ssRNAs (final concentration 0.25 μM) were determined by EMSA, ☆ represents the complex precipitation in the gel.

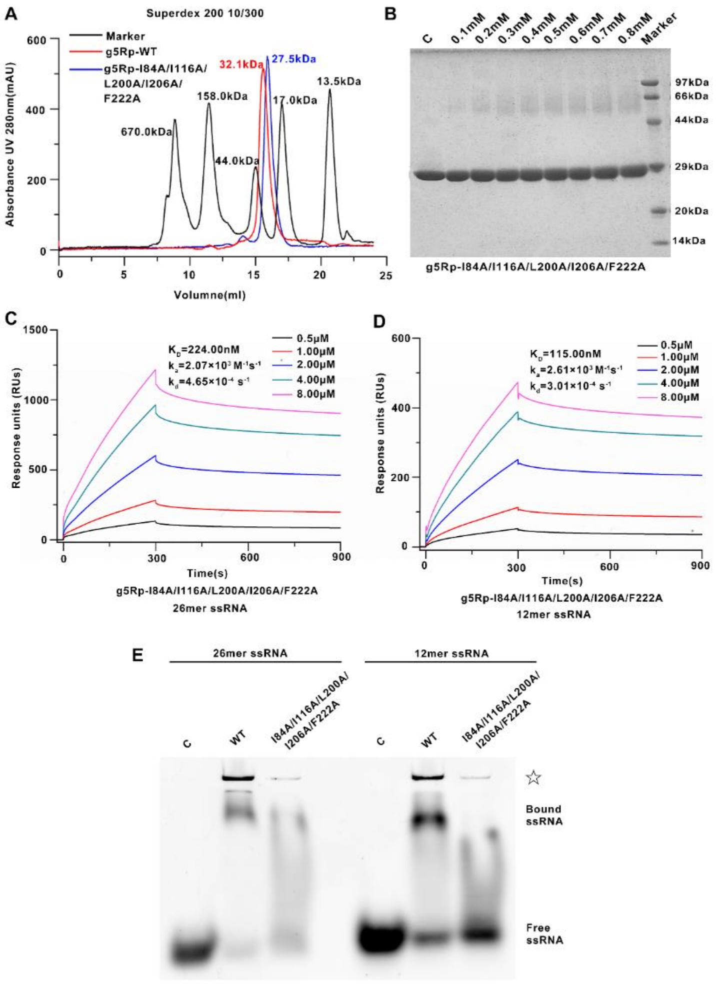

**Supplementary figure 5 Structural analysis and superposition g5Rp-InsP$_6$ complex with g5Rp.** (**A**) Surface charge distribution of g5Rp. The range of electrostatic surface potential is shown from -74.712kT/e in red color to +74.712 kT/e in blue color. InsP$_6$ is shown as sticks model. (**B**) Superposition of g5Rp with g5Rp-InsP$_6$ complex. The conformational changes are colored in cyan and red, respectively.

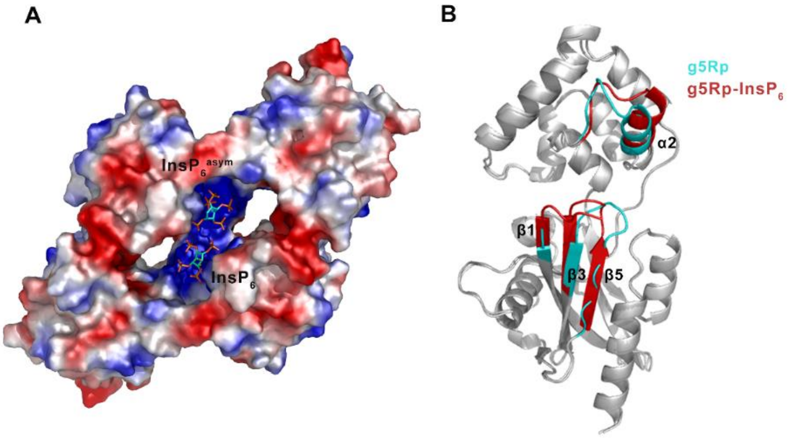

**Supplementary figure 6 The binding ability between a series of g5Rp mutants with $InsP_6$ measured by MST.** (**A-F**) MST analysis of $InsP_6$ binding to g5Rp mutants. The dissociation constant between six g5Rp mutants (Q6A, K8A, K94A, K98A, K133A, Q6K/K8A/K94A/K98A/K133A) with $InsP_6$ were calculated from three independent replicates (shown as mean±s.d.).

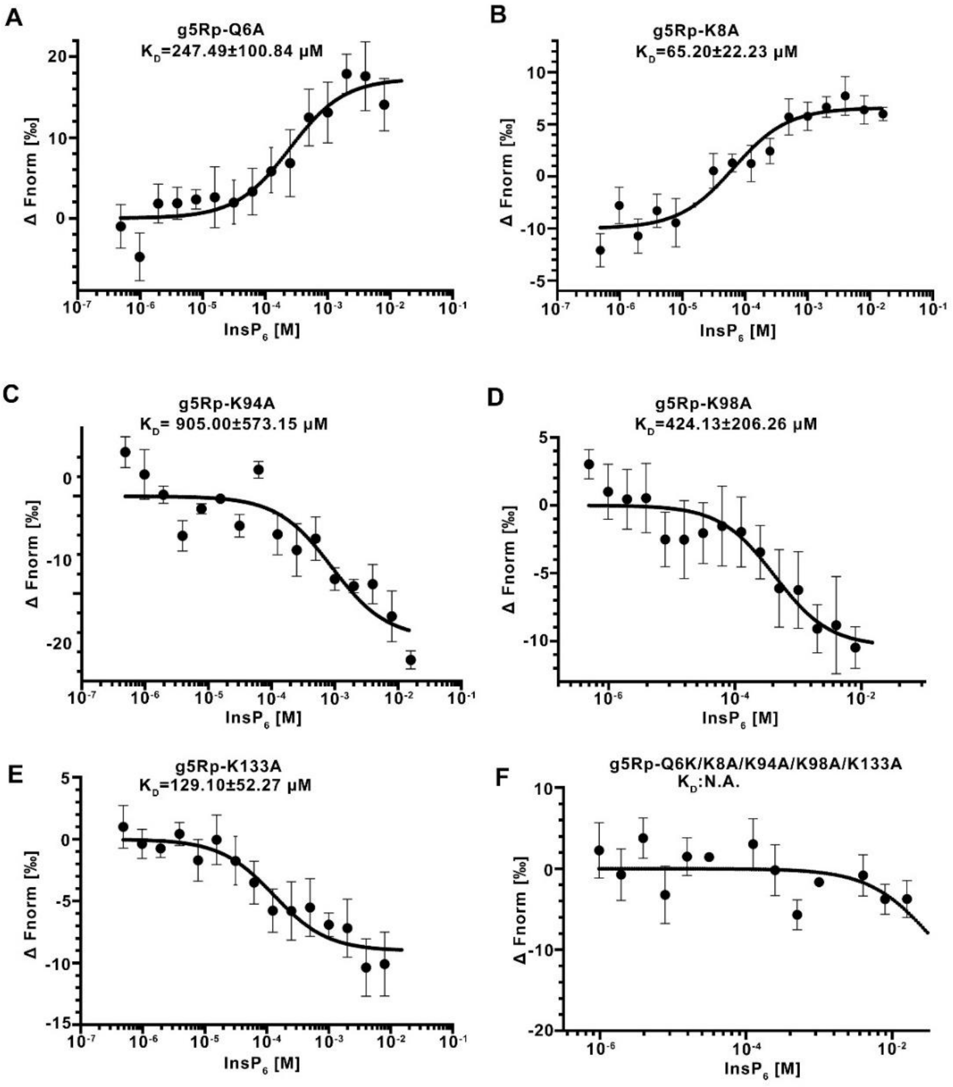

**Supplementary figure 7 Superposition and sequence alignment of the African swine fever virus decapping enzyme g5Rp with its homologs.** (**A–C**) Superposition of g5Rp with Ap4A hydrolase in complex with ATP (Aquifex aeolicus Vf5, PDB: 3I7V), Nudix hydrolase DR1025 in complex with phosphoaminophosphonic acid-guanylate ester (Deinococcus radiodurans, PDB: 1SZ3), and MTH1 in complex with 8-oxo-dGTP (Mus musculus, PDB: 5MZE). Ap4A hydrolase is shown in slate, Nudix hydrolase DR1025 is shown in marine, and MTH1 is shown in white. ATP, 8-oxo-dGTP, and GNP are shown as stick models. (**D**) Sequence alignment of g5Rp, Ap4A hydrolase, Nudix hydrolase DR1025, and MTH1. The conversed Nudix motif is marked by the black box. The conversed amino acids that bind to the substrates (ATP, 8-oxo-dGTP, and GNP) are marked by red triangles.

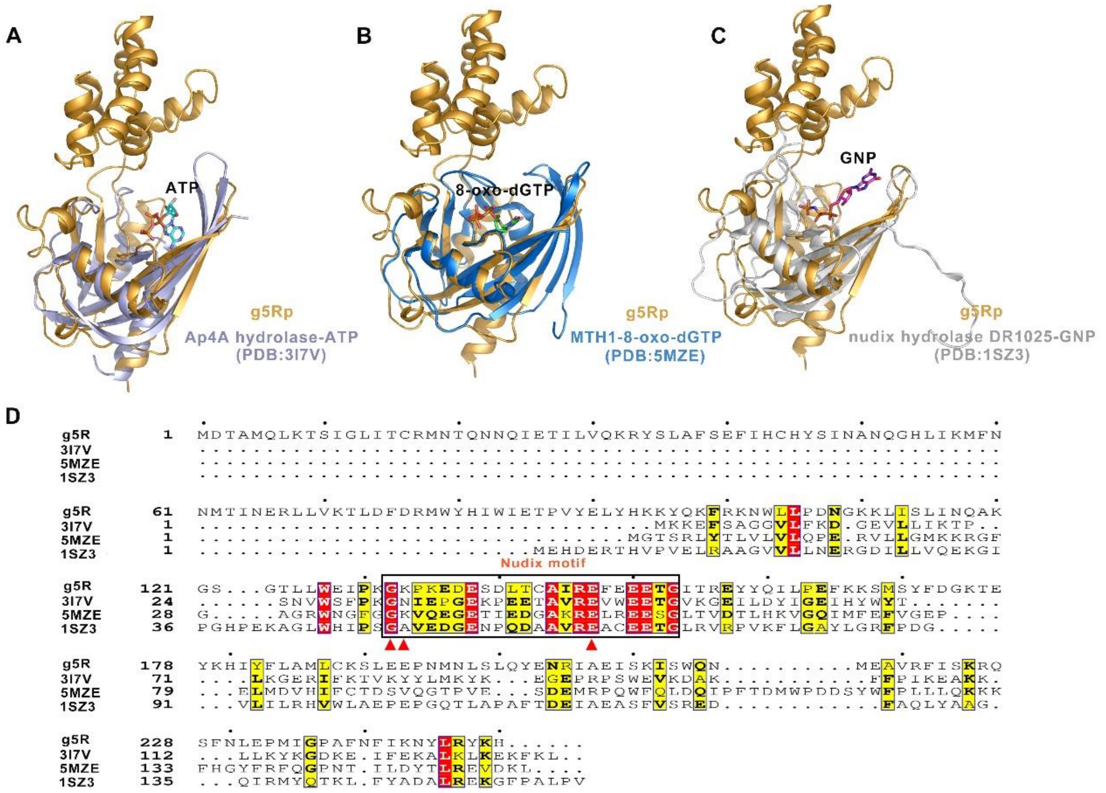

**Supplementary figure 8 The decapping activity and RNA binding abilities of g5Rp mutants.** (**A**) The comparison in binding abilities of g5Rp and Q6K/K8A/K94A/K98A/K133A (final concentration 2.00 μM) with 26mer, 12mer-ssRNAs (final concentration 0.25 μM). (**B**) The binding abilities of K8A/K131A/K133A/K135A and R221A/K225A/R226A/K243A/R247A (final concentration 2.00 μM) with 12mer, 26mer-ssRNAs (final concentration 0.25 μM). (**C**) and (**E**) The decapping activities of g5Rp mutants (Q6K/K8A/K94A/K98A/K133A, K8A/K131A/K133A/K135A and R221A/K225A/R226A/K243A/R247A). (**D**) and (**F**) The Semi-quantitative of $m^7GDP$ by graphpad prism8. (mean ± SD, $n \geq 3$, $*P < 0.05$, $**P < 0.01$, unpaired t test).

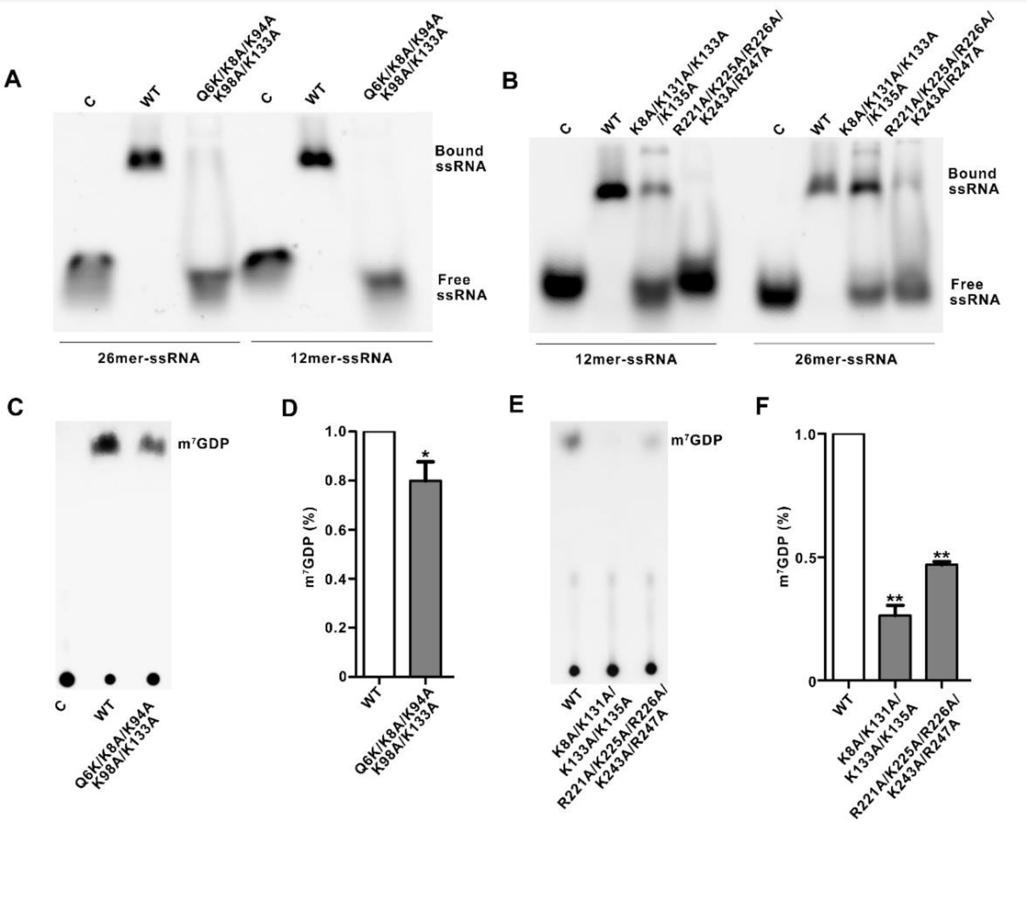

**Supplementary figure 9 Superposition of g5Rp-InsP$_6$ with DIPP1-InsP$_7$ and the thermal parameter(B-factor) distribution in g5Rp and g5Rp-InsP$_6$ complex.** (**A**) Superposition of g5Rp-InsP$_6$ with DIPP1-InsP$_7$(Homo sapiens, PDB:6PCL), g5Rp is shown in palecyan, MTH1 is shown in marine. InsP$_6$ and InsP$_7$ are shown as stick models. (**B**) B-factor distribution in Apo-g5Rp and g5Rp-InsP$_6$, shown as implemented by PyMOL. The Cα B-factors are depicted on the structure in dark blue (lowest B-factor) through to red (highest B-factor), with the radius of the ribbon increasing from low to high B-factor. The more lower B-factor is observed in the overall structure of g5Rp-InsP$_6$, with the InsP$_6$-binding sites also displaying lower than average B-factors, consistent with the InsP$_6$ contacts stabilising this region of g5Rp relative to the overall structure.

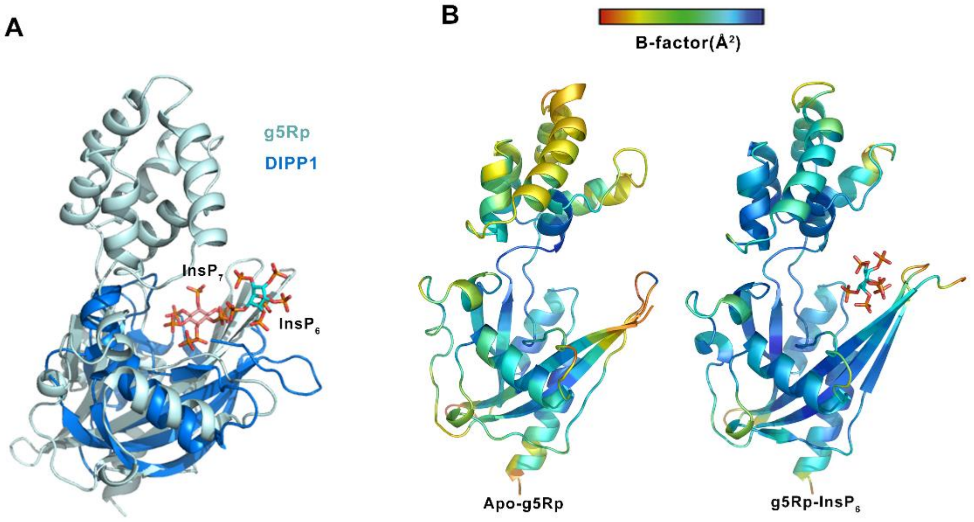

**Supplementary figure 10 The crystallization, X-ray diffraction data of g5Rp and CD spectrum of g5Rp proteins.** (**A-C**) A series of g5Rp mutants and these two g5Rp truncations secondary structure were analyzed by circular dichroism (CD) spectroscopy (**D**) The apo se-g5Rp crystal. (**E**) The g5Rp-$InsP_6$ complex crystal. (**F**) X-Ray diffraction point atlas of apo se-g5Rp. (**G**) X-Ray diffraction point atlas of the g5Rp-$InsP_6$ complex.

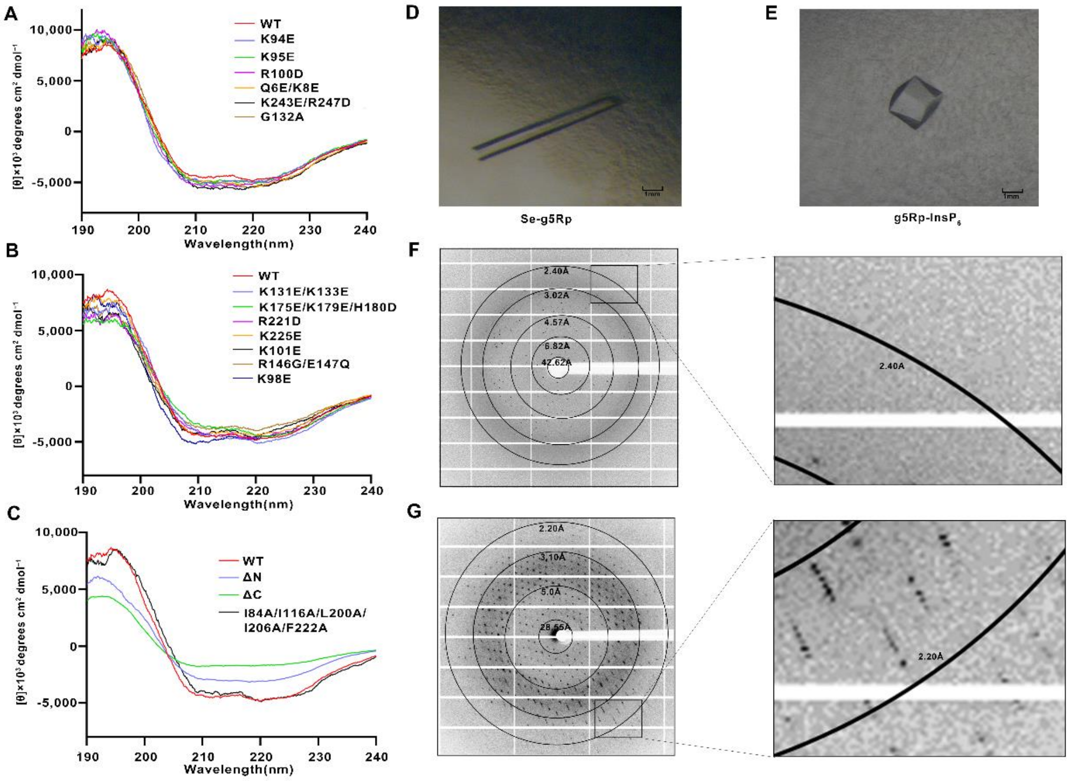

### Supplementary tables

#### Supplementary table 1. Kinetic analysis of SPR

| ligand | analyte | KD | ka (1/Ms) | kd (1/s) | Rmax (RU) | $Chi^2(RU^2)$ |
|---|---|---|---|---|---|---|
| 26mer RNA | g5Rp-WT | 4.48E-08 | 1.14e+04 | 5.10e-04 | 38.8 | 2.00 |
| | | 3.81E-08 | 1.61e+04 | 6.12e-04 | 33.5 | 3.92 |
| | G5Rp-△C | 5.07E-08 | 1.85e+04 | 9.38e-04 | 63.2 | 10.4 |
| | | 4.70E-08 | 1.47e+04 | 6.92e-04 | 46.1 | 4.04 |
| | G5Rp-I84A/I116A/ | 2.24E-07 | 2.07e+03 | 4.65e-04 | 1142.7 | 1930 |
| | L200A/I206A/F206A | 5.75E-07 | 1.05e+03 | 6.05e-04 | 1013.0 | 1470 |
| 12mer RNA | g5Rp-WT | 1.64E-07 | 3.20e+03 | 5.25e-04 | 112.9 | 1.23 |
| | | 1.15E-07 | 5.33e+03 | 6.13e-04 | 77.8 | 4.51 |
| | G5Rp-△C | 3.90E-08 | 2.12e+04 | 8.28e-04 | 62.3 | 11.6 |
| | | 2.87E-08 | 2.84e+04 | 8.14e-04 | 46.0 | 5.20 |
| | G5Rp-I84A/I116A/ | 1.15E-07 | 2.61e+03 | 3.01e-04 | 435.1 | 324 |
| | L200A/I206A/F206A | 2.25E-07 | 2.52e+03 | 5.68e-04 | 348.2 | 136 |

#### **Supplementary table 2.** Data from the Dali server

| PDB | Protein | Organism(s) | Classification | Z-Scores |
|---|---|---|---|---|
| 5CFJ | PF3D7_0520600 | *Plasmodium falciparum* (isolate 3D7) | Asymmetrical diadenosine tetraphosphate hydrolase | 17.9 |
| 1KTG | ndx-4 | *Caenorhabditis elegans* | Asymmetrical diadenosine tetraphosphate hydrolase | 17.3 |
| 1XSB | -NUDT2 | *Homo sapiens* | Asymmetrical diadenosine tetraphosphate hydrolase | 16.5 |
| 4MPO | -CTL0140 | *Chlamydia trachomatis* serovar L2 | Asymmetrical diadenosine tetraphosphate hydrolase | 16.5 |
| 3I7V | Ap4A hydrolase | *Aquifex aeolicus* (strain VF5) | Asymmetrical diadenosine tetraphosphate hydrolase | 15.5 |
| 1VC8 | Ndx1 | *Thermus thermophilus* | Diadenosine hexaphosphate hydrolase | 15.3 |
| 3GZ8 | NrtR | *Shewanella oneidensis* (strain MR-1) | Transcriptional repressor of nicotinamide riboside | 14.8 |
| 1SZ3 | DR1025 | *Deinococcus radiodurans* | MutT/Nudix family protein | 14.8 |
| 5GG7 | MutT1 | *Mycolicibacterium smegmatis* | 8-Oxo-(d)GTP phosphatase | 14.7 |
| 5KQ4 | DCP2 | *Homo sapiens, Schizosaccharomyces pombe* | mRNA-decapping enzyme | 14.6 |

| | | | | |
|---|---|---|---|---|
| 6VCM | RppH | *Escherichia co*li (strain K12) | RNA pyrophosphohydrolase | 14.2 |
| 2QJO | Slr0787 | *Synechocystis* sp. | Bifunctional NMN adenylyltransferase/Nudix hydrolase | 14.0 |
| 6AM0 | DCP2 | *Kluyveromyces lactis* | mRNA-decapping enzyme | 13.8 |
| 3I9X | Lin0490 | *Listeria innocua* serovar 6a | MutT/Nudix family protein | 13.7 |
| 2FML | EF1141 | *Enterococcus faecalis* | MutT/Nudix family protein | 13.3 |
| 5DEQ | nagR | *Bacteroides thetaiotaomicron* | GntR family transcriptional regulator | 12.9 |
| 1KHZ | - nudF | *Escherichia coli* | ADP-ribose pyrophosphatase | 12.9 |
| 6FL4 | NUDT1 | *Arabidopsis thaliana* | Nudix hydrolase | 12.7 |
| 4ZBP | NUDT7 | *Arabidopsis thaliana* | MutT/Nudix family protein | 12.7 |
| 2O5W | Nudb | *Escherichia coli* | Nudix NTP hydrolase | 12.5 |
| 3Q4I | BC2032 | *Bacillus cereus* | Phosphohydrolase (MutT/Nudix family protein) | 12.4 |
| 2A8R | Nudt16 | *Xenopus laevis* | U8 snoRNA-decapping enzyme | 12.4 |
| 5C7T | NudF | *Bdellovibrio bacteriovorus* | ADP-ribose pyrophosphatase | 12.3 |
| 1JKN | Ap4A | *Lupinus angustifolius* | Nudix enzyme diadenosine tetraphosphate hydrolase | 12.3 |
| 6U9X | MERS1 | *Trypanosoma brucei* | Mitochondrial edited mRNA stability factor | 12.2 |
| 6DT3 | NudI | *Klebsiella pneumoniae* | Nucleoside triphosphatase | 12.0 |
| 5IW4 | Nudc | *Escherichia coli* | NADH pyrophosphatase | 12.0 |
| 6O3P | Nudt12 | *Mus musculus* | Peroxisomal NADH pyrophosphatase | 11.8 |
| 5OTN | MTH1 | *Danio rerio* | Nudix nucleoside diphosphate | 11.8 |
| 3A6V | MutT | *Escherichia coli* | 8-Oxo-(d)GTP phosphatase | 11.7 |
| 6SCX | NUDT12 | *Homo sapiens* | Peroxisomal NADH pyrophosphatase | 11.6 |
| 3EF5 | MutT | *Bdellovibrio bacteriovorus* | Probable pyrophosphohydrolase | 11.6 |
| 5ZRO | MutT2 | *Mycolicibacterium smegmatis* | Nudix hydrolase | 11.5 |
| 5MZE | MTH1 | *Mus musculus* | 7,8-Dihydro-8-oxoguanine triphosphatase | 11.1 |
| 1V8L | Ndx4 | *Thermus thermophilus* | ADP-ribose pyrophosphatase | 11.1 |
| 5Z78 | NUDT16L1 | *Homo sapiens, Mus musculus* | Tudor-interacting repair regulator protein | 11.0 |
| 3WHW | NUDT1 | *Homo sapiens* | 7,8-Dihydro-8-oxoguanine triphosphatase | 11.0 |
| 1MQW | Rv1700 | *Mycobacterium tuberculosis* | ADP-ribose pyrophosphatase | 10.8 |
| 4YOQ | MutY | Synthetic construct, *Geobacillus stearothermophilus* | Adenine DNA glycosylase | 10.3 |

| | | | | |
|---|---|---|---|---|
| 1RRQ | MutY | *Geobacillus stearothermophilus* | Hydrolase/DNA | 10.1 |
| 3MDI | NUDT21 | *Homo sapiens* | Cleavage and polyadenylation specificity factor | 10.0 |
| 3G0Q | MutY | *Geobacillus stearothermophilus* | Adenine DNA glycosylase | 10.0 |
| 2I6K | IDI1 | *Homo sapiens* | Isopentenyl-diphosphate delta-isomerase | 9.8 |
| 1PVF | Idi | *Escherichia coli* | Isopentenyl-diphosphate delta-isomerase | 9.8 |
| 4DX8 | KRIT1 | *Homo sapiens* | Krev interaction trapped protein | 9.2 |
| 6PUS | TRPM2 | *Homo sapie*ns | Transient receptor potential cation channel | 8.9 |

**Supplementary table 3.** Primer sequences for generation of the g5Rp mutants

| Primer name | Sense sequence（5′–3′） | Orientation |
|---|---|---|
| R146G/E147Q-F | ATCTGACCTGTGCAATTGGCCAATTTGA | Forward |
| R146G/E147Q -R | GGCCAATTGCACAGGTCAGATCGCTTTC | Reverse |
| Q6E/K8E-F | GAGCTGGAAACCAGTATTGGTCTGATTACCTGCCGT | Forward |
| Q6E/K8E -R | CCAATACTGGTTTCCAGCTCCATGGCGGTATCCAT | Reverse |
| K131E/K133E-F | GAAATTCCGGAAGGTGAACCGAAAGAAGATGAAA | Forward |
| K131E/K133E -R | CACCTTCCGGAATTTCCCACAGCAGGGTGCCACT | Reverse |
| K175E/K179E/H180E-F | GTGAAACCGAATATGAAGACATCTATTTCCTGGCCA | Forward |
| K175E/K179E/H180E -R | GTCTTCATATTCGGTTTCACCGTCAAAATAGCTCAT | Reverse |
| K94E-F | GTTTATGAACTGTATCATGAAAAAATACCAGAAGT | Forward |
| K94E -R | CATGATACAGTTCATAAACCGGGGTTTCAATCC | Reverse |
| K95E -F | TATGAACTGTATCATAAAGAATACCAGAAGTTCC | Forward |
| K95E -R | CTTTATGATACAGTTCATAAACCGGGGTTTCAA | Reverse |
| K98E -F | TATCATAAAAAATACCAGGAGTTCCGCAAAAATT | Forward |
| K98E -R | CCTGGTATTTTTTATGATACAGTTCATAAACCGG | Reverse |
| R100D-F | AAAAAATACCAGAAGTTCGACAAAAATTGGCTGCT | Forward |
| R100D-R | TCGAACTTCTGGTATTTTTTATGATACAGTTCATA | Reverse |
| K101E-F | AAATACCAGAAGTTCCGCGAAAATTGGCTGCTGC | Forward |
| K101E-R | CGCGGAACTTCTGGTATTTTTTATGATACAGTTC | Reverse |
| R221D-F | CAGAATATGGAAGCCGTTGATTTTATTAGCAAACG | Forward |
| R221D-R | TCAACGGCTTCCATATTCTGCCAACTAATTTTGCT | Reverse |
| K225E-F | GCCGTTCGTTTTATTAGCGAACGTCAGAGCTTTA | Forward |
| K225E-R | CGCTAATAAAACGAACGGCTTCCATATTCTGCCA | Reverse |
| K243E/R247D-F | TATTGAGAATTATCTGGACTACAAGCATTAA | Forward |
| K243E/R247D -R | TCCAGATAATTCTCAATAAAATTGAATGCCGGACCA | Reverse |
| g5Rp-△N-F | GTGCAGAAACGTTATAGTCTGCTGTGGGAAATTCCG | Forward |

| | | |
|---|---|---|
| g5Rp-△N-R | CGGAATTTCCCACAGCAGACTATAACGTTTCTGCAC | Reverse |
| g5Rp-△C-F | CTGGTGCCGCGCGGCAGCCTGGCATTTTCAGAATTC | Forward |
| g5Rp-△C-R | GATGATGATTCCTCCTCCGGTGCCACTACCTTTTGC | Reverse |
| G132A-F | CCGAAAGCTAAACCGAAAGAAGATGAAAGCGATCTG | Forward |
| G132A-R | TTTCGGTTTAGCTTTCGGAATTTCCCACAGCAGGGT | Reverse |
| K133A-F | GAAAGGTGCACCGAAAGAAGATGAAAGCGATCTGAC | Forward |
| K133A-R | TCTTTCGGTGCACCTTTCGGAATTTCCCACAGCAGG | Reverse |
| Q6A-F | GCGCTGAAAACCAGTATTGGTCTGATTACCTGC | Forward |
| Q6A-R | AATACTGGTTTTCAGCGCCATGGCGGTATCCAT | Reverse |
| K8A-F | CTGGCAACCAGTATTGGTCTGATTACCTGCCGTATG | Forward |
| K8A-R | ACCAATACTGGTTGCCAGCTGCATGGCGGTATCCAT | Reverse |
| K94A-F | TTATGAACTGTATCATGCAAAATACCAGAAGTTCCG | Forward |
| K94A-R | GCATGATACAGTTCATAAACCGGGGTTTCAATCCAA | Reverse |
| K131A/K133A/K135A-F | GCAGGTGCACCGGCAGAAGATGAAAGCGATCTGACC | Forward |
| K131A/K133A/K135A-R | TTCTGCCGGTGCACCTGCCGGAATTTCCCACAGCAG | Reverse |
| R221A/K225A/R226A-F | TGCTTTTATTAGCGCAGCTCAGAGCTTTAATCTGGA | Forward |
| R221A/K225A/R226A-R | GCTGCGCTAATAAAAGCAACGGCTTCCATATTCTGC | Reverse |
| K243A/R247A-F | TATTGCGAATTATCTGGCCTACAAGCATTAA | Forward |
| K243A/R247A-R | GCCAGATAATTCGCAATAAAATTGAATGCCGGACCA | Reverse |
| I84A-F | TATTTGGGCTGAAACCCCGGTTTATGAACTGTAT | Forward |
| I84A-R | GGGGTTTCAGCCCAAATATGATACCACATGCGAT | Reverse |
| I116A-F | TAGTCTGGCCAACCAGGCAAAAGGTAGTGGCA | Forward |
| I116A-R | GCCTGGTTGGCCAGACTAATCAGTTTTTTACCAT | Reverse |
| L200A-F | ATATGAATCTGAGTGCGCAGTATGAAAATCGC | Forward |
| L200A-R | GCGCACTCAGATTCATATTCGGTTCTTCCAGGCT | Reverse |
| I206A-F | GCTGCCGAAATTAGCAAAATTAGTTGGCAGAATA | Forward |
| I206A-R | TTTGCTAATTTCGGCAGCGCGATTTTCATACTGC | Reverse |
| F222A-F | CGTTCGTGCTATTAGCAAACGTCAGAGCTTTA | Forward |
| F222A-R | TTGCTAATAGCACGAACGGCTTCCATATTCTGCC | Reverse |
